# Unobserved Sequence Space Has Many Functional Proteins

**DOI:** 10.64898/2026.08.24.746828

**Authors:** Nathan Diplock, Thomas Desautels, Riyaz Haque, Micaela Finney, Daniel J. Wackelin, Christina S. Kang-Yun, Dan M. Park, Sean P. Leonard

## Abstract

The distribution of functional proteins across amino acid sequence space, and the proportion of functional space covered by existing proteins, remains unknown^1,2^. Illuminating this distribution is integral to understanding protein evolution and advancing protein design. The recent explosion of AI/ML protein design tools presents an opportunity to explore protein sequence space distant from extant proteins, but these tools remain poorly validated. Here, we determine that portions of protein sequence space, despite being unobserved in nature, contains many functional proteins that cannot be predicted accurately *in silico*. We measure experimental fitness of highly diverse proteins across 3 families and assemble the largest known dataset of diverse, functionally labeled natural protein orthologs and new-to-nature proteins. For each family, we observe many functional, new-to-nature sequences with low amino acid identity to existing orthologs. Sequence-based scoring metrics, especially Potts models and protein language models, provide accurate but inconsistent and highly correlated function predictions. Empirical protein fitness landscapes are rugged, and predictions of function do not consistently capture either the local shape or global trends of the empirical fitness landscapes. Finally, we find extensive functional sequence space between existing proteins in each family, providing experimental support for the hypothesis that natural protein sequences explored by evolution represent a miniscule fraction of all possible functional sequences.

## Introduction

Over 50 years ago, John Maynard Smith questioned the extent to which all functional protein sequence space is occupied by observed natural protein sequences, and this has remained an open question in biology^1^. Answering this question is essential for understanding protein evolution as well as advancing protein design and requires characterizing the distribution of function across protein sequence space. Determining the distribution of function in protein sequence space relative to existing natural sequences in “occupied” sequence space is complicated by both the expansiveness of “complete” protein sequence space, with possible combinations equaling 20^N^, where N is protein length, and the context-specific nature of how function is defined. Occupied sequence space containing all natural proteins may represent a miniscule fraction of functional plausible sequence space, although the distribution of function in the remaining fraction of “unobserved” sequence space has not been characterized.

The distribution of protein function can be visualized as a protein fitness landscape. Current empirical protein fitness landscapes come from local explorations using deep mutational scanning (DMS) of dense, combinatorial mutations of up to 4 positions in a single protein^3–7^, a small fraction of the sequence space occupied by existing proteins, and an even smaller fraction of the unobserved sequence space that exists between known proteins. Numerous examples exist of small mutational steps that lead to improved or novel function of existing proteins^3–5,8^, but the distribution of functional sequences and the shape of protein fitness landscapes across diverse, plausible sequence space is unexplored^2^. One dataset evaluating several thousand diverse natural orthologs and unobserved sequences exists for the chorismite mutase protein family^9^, but unobserved sequence space was sampled to maximize predicted function and tested new-to-nature sequences were similar to existing natural orthologs as a result. In addition, while some large-scale comparisons of fitness of natural proteins exist^10–12^, functional validation of natural sequences remains sparse, preventing exploration of broader protein family fitness landscapes. While many theories exist on the shape of the distribution of function in sequence space, from isolated islands of function, to narrow traversable ridges, empirical protein fitness landscapes comprised of diverse proteins in occupied and unobserved sequence space have never been explored at scale^13^.

Recent advancements in AI/ML protein design tools have enabled generative protein design into unobserved sequence space^9,14–16^ and these tools can be used to predict the function of protein sequences, with the potential for accurate generalization to new-to-nature proteins^14^. However, it remains unclear if AI/ML models can accurately predict the function of proteins in unobserved sequence space, as these generally exist outside of established training data for these algorithms^17–20^. Accurate prediction into unobserved sequence space would enable *in-silico* evaluation of diverse sequence space across many protein families.

Several attempts have been made to benchmark the accuracy of protein function prediction using a variety of models and metrics^14,21,22^, but these benchmarking datasets do not contain large-scale, functionally validated, and diverse new-to-nature sequences. DMS benchmarking datasets like those within ProteinGym are often used to interrogate variant effect prediction, but they do not contain functionally labeled variants that are distant from occupied sequence space. Moreover, while often an assumption made during model training, the protein sequences and/or structures that are used to train AI/ML protein models are likely not all functional in their native context, and certainly not all functional in the context of a given protein design task^9,10^. Data evaluating the extent of context-specific functionality for sequences within and beyond underlying model distributions will help advance protein design through insight into the distribution of protein function in training data and the relationship between sequence and function^23^. Overall, the ability of protein AI/ML models to capture function across proteins in the underlying model distribution and generalize into unobserved sequence space is not known.

We set out to determine if functional sequence space commonly exists outside of the sequence space occupied by natural proteins and to explore the accuracy of protein scoring metrics across occupied and unobserved sequence space. We generate functional new-to-nature sequences well outside of natural sequence space using family-specific Potts models, an approach previously demonstrated to produce functional sequences distant from existing proteins^9^. We create the largest known library of diverse, new-to-nature and natural proteins with experimentally determined fitness scores across 3 distinct protein families, comprised of 3,000 natural orthologs and 10,000 proteins in unobserved sequence space. We use pooled growth-based assays to determine protein function and leverage this dataset to explore predictors of protein function across real protein fitness landscapes. We show that protein function prediction fails to capture the local resolution of protein fitness landscapes, and that extensive unobserved functional sequence space exists across three protein families. Our labeled datasets are made available to advance model training and benchmarking across the protein design community.

## Results

### Diverse orthologs complement three conditionally essential *E. coli* genes

We first identified candidate genes suitable for high-throughput characterization of protein function by testing a subset of conditionally essential *E. coli* genes that have simple non-permissive growth conditions. In these assays, an *E. coli* deletion mutant strain is complemented with a functional or non-functional protein, linking cell survival to protein function. We tested KEIO^24^ knockouts for 19 candidate genes and selected three genes that encode for proteins with diverse functions and robust growth-based assays: 1) SdhE, an assembly factor required for flavinylation of SdhA and downstream succinate metabolism^25^; 2) CrcB, an inner membrane fluoride ion export channel required for growth in media containing fluoride^26^; and 3) PanD, an aspartate decarboxylase that produces β-alanine, a precursor to Coenzyme A and a requirement for growth in minimal media^27^ (Fig. 1A). For each gene the *E. coli* knockout strains displayed expected conditionally essential growth in non-permissive media conditions (see Methods), and that growth could be restored by complementation of the corresponding *E. coli* wild-type (WT) gene as well as a set of diverse orthologs (Fig. 1B).

**Figure 1.**
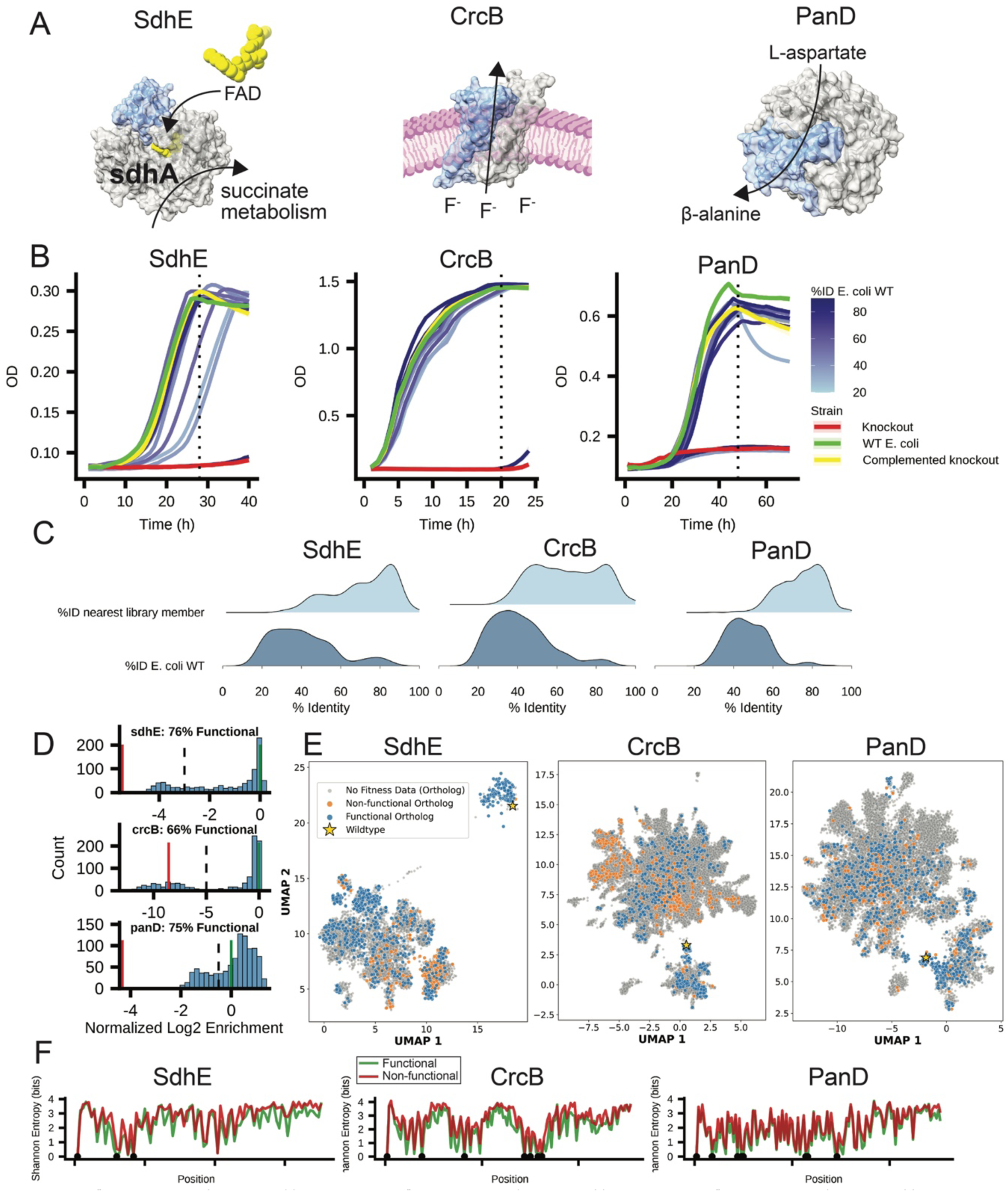
Diverse natural orthologs functionally complement conditionally essential genes in *E. coli*. (A) Three selected protein families: SdhE (left PDB:6C12) promotes flavinylation of SdhA and succinate metabolism, CrcB (center) is a fluoride efflux channel, and PanD (right) is an aspartate decarboxylase. CrcB and PanD structures were generated with AlphaFold2^19^. (B) *E. coli* knockout strains result in conditionally essential growth in non-permissive growth media (red) that can be complemented by the *E. coli* WT sequence (yellow) and diverse natural orthologs (blue). Dashed black line indicates the timepoint at which samples were collected in pooled library screens (C) Ortholog libraries selected for experimental evaluation have low pairwise identity to other library orthologs (top) and a range of sequence identity to the *E. coli* WT sequence (bottom) (D) Mean normalized log2 enrichment of tested orthologs in non-permissive media relative to permissive media normalized to the *E. coli* WT control (n=3). Black dashed vertical lines indicate the functional threshold for each library and was determined using growth assays of individual colonies (see Extended Data Fig. 3). (E) UMAP representations of one-hot encoded natural ortholog sequences across the three tested protein families (n=952 for SdhE, n=957 for CrcB, n=984 for PanD). (F) Shannon entropy across aligned functional and non-functional tested orthologs with gap. Black dots above the x-axis indicate Shannon entropy <0.1.

To define the functional ortholog landscape of SdhE, CrcB, and PanD, we first searched Uniref90^28^ and MGnigy^29^ protein databases for orthologs of the *E. coli* WT protein. We retained sequences with greater than 70% alignment coverage and less than 90% sequence identity to the original *E. coli* WT protein (Extended Data Fig. 1). To thoroughly explore the fitness of natural sequence space, we selected and synthesized 1000 diverse orthologs for each gene using stratified sampling across bins of percent identity to the *E. coli* sequence (Fig. 1C, Extended Data Fig. 1). The selected variant sequences have both low percent identity to the *E. coli* WT protein sequence, and low pairwise identity to the nearest library member (Fig. 1C). Next, we performed triplicate pooled assays by growing *E. coli* libraries containing all orthologs in permissive and non-permissive conditions, including *E. coli* WT and a non-functional ortholog spiked in as positive and negative controls. Illumina sequencing allowed us to estimate the frequency of each variant in each condition and the enrichment of functional proteins in selective media. Log2 enrichments normalized to the *E. coli* WT resulted in a clear bimodal distribution of functional and non-functional sequences, with strong agreement observed between replicates (Fig. 1D, Extended Data Fig. 2). We further validated the ortholog library-based screen and functional threshold by randomly picking 92 colonies to sequence and test through individual growth assays. Using this individual growth data, we set conservative log2 enrichment thresholds for function, allowing some functional proteins in individual growth assays to be considered non-functional in the library context but not vice-versa (Fig. 1D, Extended Data Fig. 3). Across all three genes in our panel greater than 2/3 of selected orthologs were functional (Fig. 1D).

To visualize natural sequence space, we generated UMAP plots of one-hot sequence encodings for each variant, and overlayed ortholog function data (Fig. 1E). Distinct groups of functional and non-functional sequences exist across CrcB sequence space, but not SdhE or PanD sequence space (Fig. 1E). To further examine sequence conservation across our ortholog pools, we plotted Shannon entropy for each amino acid position across aligned orthologs. Non-functional sequences tended to have slightly higher entropy than functional sequences, especially at low entropy conserved residues. Functional orthologs maintain key catalytic residues in PanD (S25, T57, Y58), a key residue for SdhE flavinylation of SdhA (R14), and key residues required for CrcB interaction with a required Na^+^ (G75, T78) (Fig. 1E). While many non-functional orthologs maintain these conserved residues, proteins lacking these conserved sites are almost always non-functional (Fig. 1E). Although key positions were highly conserved, we identified functional orthologs with 13.3%, 18.3% and 27.4% amino acid identity to the SdhE, CrcB, and PanD *E. coli* WT sequence respectively.

### Potts models built from natural orthologs regularly design functional proteins

To explore the fitness of unobserved sequences within our three protein families, we built Potts models using all natural ortholog sequences in our MSAs. As expected, the Potts models captured extensive pairwise couplings (Fig. 2A, Extended Data Fig. 4). Next, we generated 30,000-70,000 sequences from Potts models and selected 2,000-6,000 sequences for each protein family for synthesis. As with the ortholog libraries, we used stratified sampling based on percent identity to the *E. coli* sequence to ensure highly diverse sequences were included (Extended Data Fig. 1). As with the ortholog libraries, we determined the threshold for function by conducting microwell plate growth assays on 92 randomly selected designs, and ultimate categorized each protein as functional or non-functional based on log2 enrichment (Fig. 2C, Extended Data Fig. 5, Extended Data Fig. 6).

**Figure 2.**
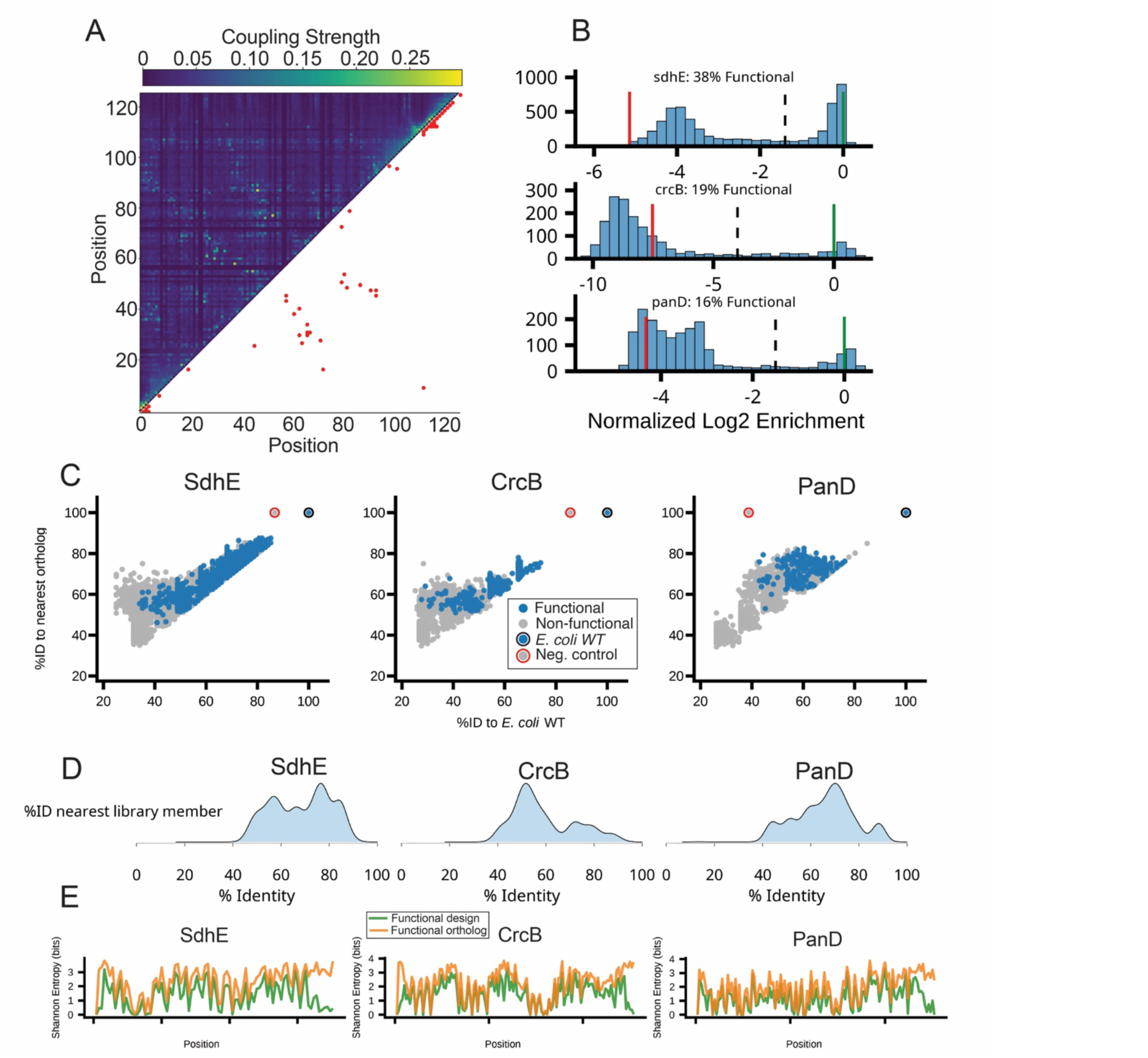
Potts models redesign functional sequences with low identity to existing natural orthologs. (A) Coupling strength of all PanD positions based on direct coupling analysis with the top 50 strongest couplings plotted in bottom right. (B) Mean normalized log2 enrichment of tested designed proteins in non-permissive media relative to permissive media normalized to the *E. coli* WT control (n=3). Black dashed lines indicate the functional threshold for each library and was determined using growth assays of individual colonies (see Extended Data Fig. 5). (C) Functional and non-functional sequences exist at low percent identity to the *E. coli* WT sequence and existing natural orthologs (n=5522 for SdhE, n=1791 for CrcB, n=1908 for PanD). (D) Selected libraries are diverse relative to existing sequences (see C) as well as relative to each other based on maximum pairwise identity within each library. (F) Shannon entropy indicates that functional orthologs generally have higher entropy than functional designs across all three protein families despite selected design and ortholog sequences having similar entropy overall (see Extended Data Fig. 7).

Simple, lightweight Potts models regularly design diverse functional sequences in unobserved sequence space for all three protein families evaluated (Fig. 2). Across our design libraries, 16% of PanD proteins, 19% of CrcB proteins, and 38% of SdhE proteins were functional (Fig. 2B). While the proportion of functional sequences decreased with the percent identity to both the *E. coli* WT sequence and existing natural sequences, functional sequences were still frequently generated with less than 60% identity to known ortholog sequences (Fig. 2C). The Shannon entropy of sequences designed using Potts models closely matched the entropy of natural orthologs, apart from the C-terminus of all proteins where new-to-nature sequences have lower entropy (Extended Data Fig. 7). This discrepancy is likely an artifact due to alignment gaps at the C-terminus of natural orthologs in our MSAs, resulting in lower diversity in these regions in the corresponding sequences sampled from Potts models. We noted that the difference in Shannon entropy between functional and non-functional designs is larger than the difference between functional and non-functional orthologs (Fig. 1F, Extended Data Fig. 7). Finally, functional orthologs generally have higher Shannon entropy than designs (Fig. 2E), suggesting that orthologs have more permissive requirements for function which Potts model designs fail to replicate.

We further confirmed the function of select proteins through heterologous and *in vitro* validation. We showed that 12 functional SdhE proteins can flavinylate SdhA *in vitro* (see Supplemental Information). Using an *in vivo* F^-^ riboswitch reporter assay, we found that CrcB function determined by library enrichment is correlated with low intracellular fluoride concentrations (see Supplemental Information). Finally, we conducted *in vitro* enzyme assays to calculate enzyme kinetics for 9 PanD proteins and showed that functional proteins determined via growth-based assays are functional aspartate decarboxylases, although only one redesigned panD variant had a higher kcat than the *E. coli* WT (see Supplemental Information).

### Simple metrics match or outperform AI predictions of new-to-nature protein function

Next, we sought to identify if existing computational tools or protein features could accurately predict the function of natural orthologs or designed protein sequences. We selected several sequence-based tools including protein family profile hidden Markov models (HMM)^30^, Potts model energy scores^31^, the autoregressive protein language model (pLM) ProGen2^32^, and the masked pLM trained for variant effect prediction ESM1v^33^. We use ESMFold^34^ generated structure predictions to calculate likelihoods from proteinMPNN^35^ and ESM inverse fold (ESM-IF)^36^ in addition to TM-scores^37^ comparing structural similarity to the *E. coli* WT. We calculated F1 scores and area under the receiver operating characteristic curve (AUC) on functional and non-functional binary fitness determined previously for each library.

Sequence-based scoring metrics accurately differentiate between functional and non-functional proteins across unobserved sequence space in design libraries, as reflected by high AUC values (Extended Data Fig. 8). However, this strong ranking performance is also observed using sequence identity alone and tempered by low F1 scores in PanD and CrcB libraries (Extended Data Fig. 8). Across natural and designed proteins, sequence-based scoring metrics generally outperform structure-based metrics. Surprisingly, relatively lightweight Potts models predict function with similar or greater accuracy than AI models across unobserved sequence space (Extended Data Fig. 8). Several scoring metrics are strongly correlated including Potts model energies and ProGen2 likelihoods, ESM1v and ProGen2 likelihoods, and ProteinMPNN and ESM-IF likelihoods (Extended Data Fig. 8, see Supplemental Information). Scoring metrics had lower accuracy when predicting the function of orthologs, compared to new-to-nature proteins, and this is likely due to tested orthologs scoring higher overall (Extended Data Fig. 8, see Supplemental Information). While several metrics are good at predicting the function across some protein families and tested libraries, no tools accurately discriminate between functional and non-functional proteins of similar sequence across any protein family or library. When a functional library protein is paired with the most sequence similar non-functional protein, no tools consistently score the functional protein more favorably than the non-functional protein (see Supplemental Information). Moreover, while pLM embeddings can cluster designed proteins by functional class more accurately than sequence distance alone, sequence distance is more accurate than pLMs for orthologs, and UMAP dimension reduction of one-hot encoded sequences can match or outperform the accuracy of pLM clustering of designs (see Supplemental Information). Taken together, our results suggest that while the evaluated protein scoring metrics can capture information about protein function for specific protein families across diverse unobserved sequence space, they are inconsistent across all protein families and inaccurate in differentiating function in local sequence space (see Supplemental information).

### Empirical protein fitness landscapes are rugged

Next, we sought to use UMAP representations of sequence space to generate and compare experimentally evaluated and predicted protein fitness landscapes. Given our observation that scoring metrics predict global fitness trends but not local fitness differences (see Supplemental Information), we hypothesized that metrics would predict smoother protein fitness landscapes with less local resolution than experimentally validated landscapes. For each protein family, we placed scaled fitness data, ProGen2 log likelihoods, or Potts model energy on the z-axis of UMAP plots containing all experimentally evaluated protein orthologs and designs. As expected, protein scoring metrics predict a smoother fitness landscape with less local resolution than ground truth experimental fitness visually and as measured qualitatively by mean landscape slope gradient and roughness calculated as mean neighbor absolute difference (Fig. 3, Extended Data Fig. 9, Extended Data Table 1). Empirical protein fitness landscapes for all three tested proteins are rugged (Fig. 3 Extended Data Fig. 9).

**Figure 3.**
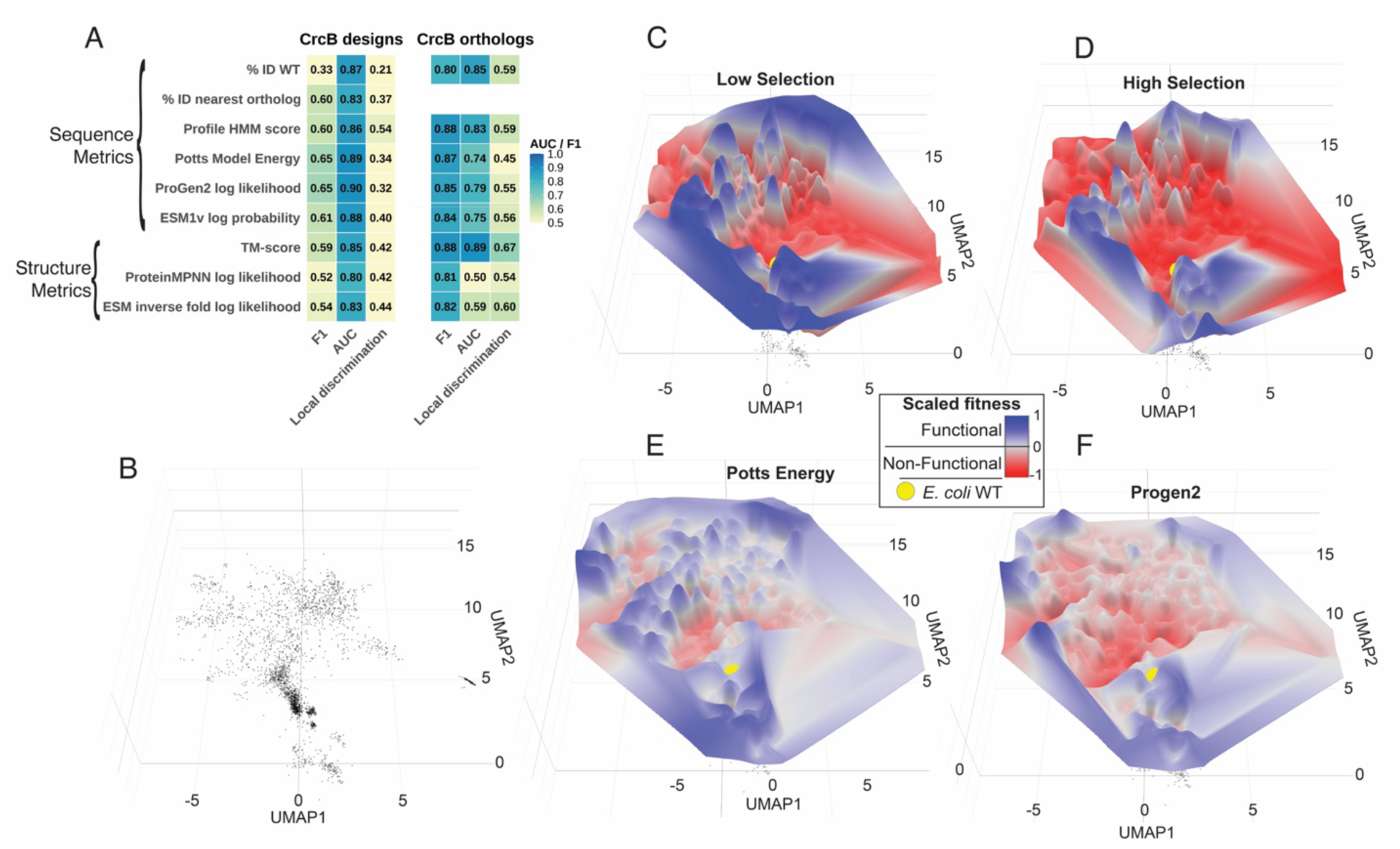
The CrcB protein fitness landscape is rugged, and no metrics accurately capture the distribution of CrcB function. (A) Accuracy of diverse sequence and structure metrics for binary function prediction across the crcB design and ortholog libraries, quantified by F1 score, AUC, and local discrimination. Local discrimination is the proportion of nearest functional/non-functional sequence pairs (based on sequence distance) where the functional protein has a more favorable score than the non-functional protein. One-hot encoded UMAP representations of tested CrcB designs orthologs on the x- and y-axis (B) are overlaid with scaled fitness values from CrcB ortholog and new-to-nature protein pooled assays under low (5mM NaF non-permissive media) selection (C) and high (50mM NaF non-permissive media) selection (D). Scaled inverted scores from Potts model energy (E) and ProGen2 (F) for all tested CrcB orthologs and new-to-nature proteins are overlayed on the z-axis of the CrcB sequence UMAP plots. The *E. coli* WT CrcB protein sequence is shown as a yellow sphere. Fitness is scaled such that 0 (white) is the estimated threshold for function across empirical and predicted landscapes (see methods).

The hypothesis that selection pressure influences the shape of fitness landscapes was first proposed nearly a century ago^38^. We sought to explore the effect of selection pressure on our experimentally evaluated landscapes. For CrcB, selection can be made more stringent by increasing fluoride concentrations. When we screened our variant pools under high selection conditions, peaks on the CrcB fitness landscape became sharper with isolated local fitness maximum compared to landscapes built with less stringent non-permissive media fitness values (Fig. 3C,D). Our data for CrcB supports the hypothesis that under strong selective pressure, it is more likely to become trapped in a local fitness maximum, while under low selective pressure, protein landscapes are more navigable. This suggests that the probability of reaching a protein global fitness maximum may be improved by using diverse starting points in sequence space^39^, especially under strong selection pressure.

### Many functional proteins exist between natural orthologs

An outstanding question in the field of biology is the extent to which extant natural protein sequences cover all possible functional sequence space^1^. While we have shown that extensive functional sequence space exists that is distinct from natural orthologs based on sequence identity (Fig. 2C), we sought to explore the relationship of unobserved functional sequence space near, distant from, and “between” natural orthologs. Since sequence space is high-dimensional, we examined relationships between designs and orthologs at the sequence level by identifying designs that were centered between two natural orthologs in sequence triplets, as well as those that were off-centered because of either high similarity to one natural ortholog or excessive distance along the paths between natural orthologs (see Supplemental information).

Across all three protein families evaluated, we found many functional protein designs between occupied sequence space (Fig. 4), and additional examples of functional sequences outside of occupied sequence space (see Supplemental information). Functional designs in unobserved sequence space are distant from existing natural orthologs and are centered in triplets, representing distant functional transitions at the sequence level between existing orthologs (Fig. 4). While most functional unobserved sequence space exists between at least one functional ortholog, we found a limited number of functional unobserved sequence space between two non-functional orthologs (Fig. 4D, bottom). To better visualize the distribution of function in unobserved sequence space, we plotted sequences through UMAP dimension reduction of one-hot sequence encodings. We identified large clusters of functional proteins in unobserved sequence space when visualized through UMAP dimension reduction (Fig. 4E-G). Overall, these results demonstrate that while occupied sequence space may adequately capture the “rules” of functionality for specific protein families, unobserved functional sequence space is expansive and designing new-to-nature sequences within the “rules” of occupied sequence space is straightforward.

**Figure 4.**
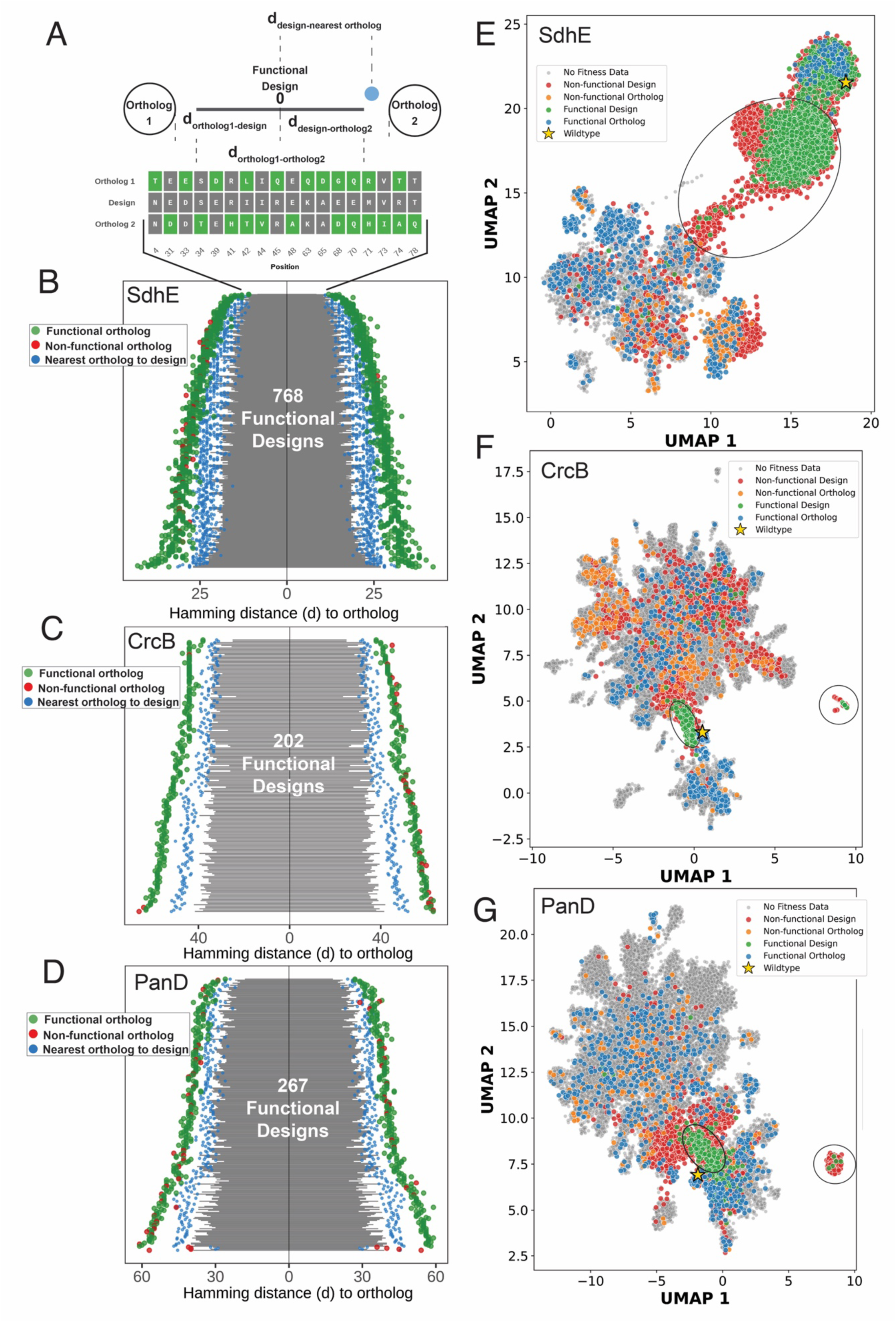
Functional sequence space exists between natural orthologs. (A) Functional new-to-nature proteins were considered as being “between” two orthologs if the designs were closer to each ortholog then each ortholog is to each other based on Hamming distance (d) in a sequence triplet. Instances where a new-to-nature protein had a Hamming distance that was 15 greater than the distance to that new-to-nature protein nearest ortholog were excluded. All variable positions for an example of the top ranked sdhE triplet shown, with green positions indicating deviations from the new-to-nature sequence. All functional (B) SdhE (C) CrcB, and (D) PanD designs that exist between two orthologs in sequence space, with each row represents a unique functional design. Sequence triplets were ranked to designs that 1) lie in a direct path between orthologs in sequence space 2) are more centered between designs and 3) have minimal distance to both orthologs (see Methods). Green and red dots indicate the Hamming distance of each functional or non-functional ortholog respectively in the triplet from the new-to-nature protein. Grey bars indicate the total distance between both orthologs, cumulative left and right of the y-axis. Blue dots indicate the distance between a new-to-nature protein and the nearest corresponding ortholog, which may not have been characterized experimentally in our assays. When present, examples of functional new-to-nature proteins between two non-functional orthologs are plotted at the bottom of each panel. UMAP plots of (E) SdhE (F) CrcB, and (G) PanD using one-hot encoded sequences for all tested designs and orthologs. Black ovals indicate approximate regions of sequence space occupied only by functional designed sequences (also see Fig. 1E).

## Discussion

In this work, we generated the largest known dataset of functionally validated, highly diverse protein sequences and used it to explore the distribution of function across protein sequence space for three protein families. Surprisingly, we found Potts models routinely generate a high proportion of functional sequences diverse from existing orthologs. While the distribution of function across sequence space can theoretically be predicted *in silico* using protein metrics, we find that no metrics can consistently predict the function of diverse orthologs or new-to-nature sequences across all three evaluated protein families. We show that diverse, unobserved sequence space contains many functional proteins across all three protein families, suggesting that functional sequence space is not adequately represented through the occupied sequence space of orthologs in a protein family. Moreover, we show that empirically fitness landscapes are rugged, and that existing tools for predicting the distribution of function across sequence space fail to capture the ruggedness of protein function for certain protein families as well as the global shape of protein fitness landscapes. Finally, we show that landscapes become less navigable with increased selection pressure, supporting a long-held theory about fitness landscapes^38^.

Our results provide additional support for findings from previous work demonstrating that large portions of occupied sequence space are non-functional, and that function exists in unobserved sequence space. Previous research using *E. coli* growth-based complementation of over 1000 chorismate mutase homologs^9^ and over 1000 phosphopantetheine adenylyltransferase homologs^10^ has shown 38% and 70% of homologs were functional respectively. Our findings that greater than 2/3 of tested orthologs are functional across three protein families supports the finding that occupied sequence space is largely functional, but many non-functional natural sequences exist when tested in the context of *E. coli*. For the goal of protein redesign in *E. coli*, models trained on occupied sequence space contain many non-functional sequences.

It is difficult to compare the absolute degree of function identified across the three protein families tested here to previous redesign efforts, given that our sampling strategy directly overrepresented sequence space distant from natural proteins. However, while more systematically explored here, our finding that many functional sequences exist which are unique from natural sequence based on identity has been supported in prior redesign efforts. For example, Potts Model based redesigns of chorismate mutases were shown to be 30% functional, with functional designs identified between 60-70% identity to occupied sequence space. Two recent investigations using protein redesign to improve function of Cas9^15^ or reverse transcriptases^16^ generated more functional variants with 86% and 74% identity to the nearest sequence identified through BLAST respectively. Clearly, functional amino acid sequences can be routinely generated with less than 80% identity to existing sequences. Patents protecting proteins often cover sequence variants 80% identity or higher to the protein of interest, so this finding may have substantial implications for biotechnology intellectual property.

While no efforts to benchmark protein prediction tools on datasets of this scale and diversity exist, poor accuracy of protein metrics predicting function and correlations between scoring metrics have been observed in prior studies. For example, Johnson et al., (2025) generated a dataset of 144 experimentally tested sequences for two separate enzymes through a variety of approaches at 70-80% identity to natural sequences and showed that inverse folding methods and protein language models provide highly correlated predictions of function^14^. Moreover, they showed that no tools provide scores that correlate well with function across tested unobserved sequence space^14^. Our findings provide further evidence that existing metrics are inconsistently accurate and often highly correlated when they are used to predict function across diverse natural and unobserved sequence space.

Although empirical protein fitness landscapes described to date are built on local sequence space generated from DMS datasets, nearly all are rugged^2,7,11,40^. Therefore, it is not surprising that our landscapes built across a larger area of sequence space are also rugged. However, despite this ruggedness we can visualize navigable paths that circumvent fitness valleys as have been hypothesized and described previously^2,6,38^. Our results provide direct experimental evidence that increasing selection pressure makes protein landscapes less navigable, as suggested in previous studies using β -lactamase and GB1^38,40,41^. Thus, while protein fitness landscapes are rugged, selection pressure can interact directly with landscape navigability and evolutionary paths^40,41^, and landscapes are navigable despite ruggedness^6,7^, perhaps even more navigable under heterogenous or inconsistent selection pressure allowing for neutral drift^40,41^. Reaching global maxima on protein fitness landscapes in an experimental setting likely requires diverse starting points and can likely be enhanced by variable selection^8,39^.

A central conflict in molecular evolution is the observation that proteins selected at random in the vast sequence space are very unlikely to be functional^42^, and yet diverse functional proteins exist across nature with seemingly intractable fitness gaps separating them^1^. Here, we have shown that many diverse sequences within unobserved sequence space produce functional proteins. However, we fail to identify functional natural or designed sequences that diverge from highly conserved positions in protein family MSAs. This failure may be due to the low diversity at conserved sites in the MSA, and therefore low diversity in sequences sampled from Potts models, but this pattern is ubiquitous when searching sequence space by sequence homology.

The hundreds of millions of existing protein sequences can be organized into just under 30,000 protein families that each share a set of highly conserved residues key for function. Sequence diversity within protein families, and therefore across occupied sequence space, largely represents an exploration of permissible backbone sequence variation instead of highly conserved residues required for function. Despite the continually increasing number of protein sequences and structures used to train the latest protein models, observed sequences represent a small set of unique mechanisms, likely due to constrains on how evolution explores sequence space^13^. While we show that function is common in distant, unobserved sequence space, we have not explored sequence space occupied by novel molecular mechanisms that are possible but unobserved in nature. It is unlikely that models trained on existing protein structures or sequences can generalize to explore novel functions or novel mechanisms without experimentally generated training data that surveys new, diverse molecular mechanisms for a specific function^43^.

## Methods

### MSA-Based Protein Redesign

#### Generation of EV-couplings designs

To generate sequences from the EVCouplings, the multi-stage EVCouplings method was applied to generate a Potts model from which sequences were sampled using a modified version of a previously described Gibbs sampling protocol^44^. EVCouplings default parameters and JackHMMER^30^ were used to retrieve and align sequences into a multiple sequence alignments against UniRef90^28^. Batch Gibbs sampling mode was used, allowing the specification of minimum sequence Hamming distance from the query sequence, MSA sequences, and between the members of the generated sequence set, as well as maximum Hamming distance from the query sequence. Batches of fixed-length sequences were generated within these constrained brackets of sequence distance from each query sequence, ranging from 25-80% sequence identity relative to the query sequences, where the minimum allowed distances within the generated sets and to the MSA sequences were fixed equal to the minimum allowed distance to the query sequence. A total of 2000, 2000, and 5800 sequences were generated, respectively, for PanD, CrcB, and SdhE.

#### Generation of CCMpred/CCMgen designs

To generate multiple sequence alignments (MSA) for CCMpred Potts model generation, pHMMER^30^ was used to search each *E. coli* candidate protein sequence against the UniRef90^28^ and MGnify^29^ protein sequence databases. Sequences with more than 90% identity to the *E. coli* candidate protein sequence or less than 70% coverage across the MSA were removed. CCMpred^45^ was used to generate two Potts Models with couplings inferred with either persistent contrastive divergence or reduced constraints based on a provided structure and a specified contact threshold of 12 Å. Specified structures were 1X6J for sdhE, AlphaFold (AF)-P0A790 for panD, and AF-P37002 for CrcB.

For each candidate gene, CCMgen was used to generate protein sequences via three methods: 1) MCMC sampling of the Potts Model generated with persistent contrastive divergence, 2) MCMC sampling of the reduced constraints Potts Model, and 3) Phylogeny based sequence sampling of the Potts Model generated with persistent contrastive divergence. Approaches 1 and 2 generated 10,000 sequences each, for all candidate genes, while approach 3 generated a proportional number of sequences relative to the original MSA. Approach 3 resulted in 6020 for SdhE, 47,427 sequences for CrcB and 18,000 sequences for PanD. Sequences generated from couplings inferred using persistent contrastive divergence for CrcB resulted in sequences with low complexity and unrealistic highly divergent sequences. Because of this, a Potts Model was regenerated for CrcB using pseudo-likelihood maximization and used to generate both MCMC and Phylogeny based sequences for CrcB. MCMC sampling of the CrcB reduced constraints Potts model was conducted using the same method for CrcB as for the other redesign candidates.

### Selection of Proteins for Experimental Assays

#### Selection of orthologs

Full length amino acid sequences for orthologs identified in the MSAs generated for CCMpred and MSAs generated by EV-couplings were combined and all sequences equal to or up to 10 amino acids less than the length of the *E. coli* WT sequence were retained. In addition, sequences that did not start with a methionine and sequences with non-standard amino acids were removed. For the initial small-scale screen of 10 orthologs for each redesign candidate gene, protein sequences were separated into 5 bins based on sequence identity to *E. coli* WT, and 2 sequences were selected from each bin. For pooled screens, sequences were also selected from the ortholog pools using stratified sampling based on percent identity to the *E. coli* WT. Sequences were separated into 20 bins across the range of percent sequence identity to *E. coli* WT in the ortholog pool, and up to 80-100 sequences were selected from each bin. Since some bins had less than the desired sample size, an additional small pool of sequences was selected randomly to reach the final 1000 sequence sample.

#### Selection of designs

For CrcB and PanD, 1000 CCMgen and 1000 EV-couplings designs were selected for experimental assays while for SdhE 3000 CCMgen and 3000 EV-couplings designs were selected for experimental assays. Designs from EV-couplings and CCMgen were selected following the same procedure for each candidate. ESMfold^34^ was used to generate structure predictions of each design, and USAlign^37^ was used to calculate structural similarity TM-scores relative to the *E. coli* WT structure. A random sample of designs with TM-scores above 0.85 was taken to achieve 25% of our total designs selected for experimental assays. The remaining 75% of the designs were selected based on the same stratified sampling approach used for ortholog selection with an additional small portion of the pools selected using random sampling to achieve the desired final number of designs of experimental assays.

### Experimental assays

#### Bacterial strains, cloning and library construction

Kanamycin resistance cassette insertions in candidate genes were transduced from Keio Collection strains into *E. coli* W3110 with P1 phage, and transduction was confirmed by PCR. SdhE orthologs and designs were ordered as separate oligo pools through Twist Biosciences and amplified with KAPA HiFi HotStart DNA Polymerase. CrcB and PanD orthologs and designs for were ordered as multiplexed gene fragments from Twist Bioscences and used directly in cloning. Both PanD pools were cloned via HiFi assembly into a pBAD33.1 vector modified to contain a medium expression ribosome binding site. All SdhE and CrcB libraries were cloned via HiFi assembly into a pBLxG2 vector modified to contain a chloramphenicol resistance gene in place of the original kanamycin resistance gene. *E. coli* WT SdhE, PanD and CrcB were synthesized and cloned into the modified pBLxG2 vector described for our libraries. *E. coli* WT SdhE, PanD and CrcB in addition to orthologs selected for small scale screening were synthesized by Integrated DNA Technologies and cloned into the pBAD33.1 vector described for our libraries via HiFi assembly. All DNA sequences were codon optimized for *E. coli* using the Twist Biosciences codon optimization tool, and all constructs were confirmed by sequencing.

All library HiFi assemblies were transformed into NEB 10-beta electrocompetent cells and the entire transformations were added to 50mL of LB with antibiotics following outgrowth. Plasmid DNA was extracted from pooled transformations and transformed into to the W3110 strain with the corresponding knockout and transferred to 50mL of LB with appropriate antibiotics following outgrowth. Colony counts of all transformations were conducted to assure that at least 30 times as many transformants as proteins in each library were obtained. In addition, whole plasmid sequencing was conducted on 5-10 single colonies from each library following transformation into NEB 10-beta.

#### Assay growth conditions

Libraries containing CrcB and PanD orthologs and designed sequences were inoculated directly from glycerol stocks into LB containing antibiotics, while SdhE ortholog and design variant libraries were inoculated into M9 glucose with antibiotics. For all libraries, a 1/1000 dilution of each library glycerol stock was inoculated into 50mL of media and cultures were grown overnight at 37°C. For each library, 3 replicate cultures grown from 3 separate glycerol stocks were inoculated. In addition, 3 replicates of the appropriate positive and negative control strains were grown separately at 37°C overnight in the same media conditions as variant libraries. After overnight growth, separate positive and negative control replicates were mixed into the corresponding library 50mL cultures at a dilution equivalent to the number of proteins in that library. For example, the SdhE design library contained 6000 proteins, and each control was added into the library at a 1/6000 dilution.

Both PanD and SdhE library cultures with added controls were pelleted and washed 2 times in M9 media with no carbon. Each washed replicate of SdhE ortholog and design libraries was added to 50mL of growth non-permissive M9 + 0.3% succinate, and growth permissive M9 + 0.3% succinate, 0.2% glucose media at a final dilution of 1/1000. Each washed replicate of PanD design variants was added to 50mL of growth non-permissive M9 + 0.4% glycerol + 0.01% arabinose and growth permissive M9 + 0.4% glycerol + 0.01% arabinose + 2mg/L ϕ3-alanine media at a final dilution of 1/1000. To minimize cross feeding from a library with a high proportion of functionality, each washed replicate of the PanD ortholog library was added to 250mL of the same media outlined for panD design libraries, at a final dilution of 1/10000.

Finally, each CrcB ortholog and design library replicate was added to 50mL growth permissive LB, and growth non-permissive LB + 5mM NaF, and LB + 50mM NaF media all containing 0.01μM inducer, at a final dilution of 1/1000. Cultures were grown at 30°C, and samples were take at 20, 28, 48, and 32 hours for CrcB libraries, SdhE libraries, the PanD design library and the PanD ortholog library respectively. Plasmid DNA was extracted from samples, and library constructions and 150 paired end Illumina sequencing were conducted by Neochromosome.

#### Microplate assays of protein variants

Growth of orthologs selected for initial small-scale screening was assayed in microplates following the same media and wash conditions outlined for PanD, SdhE and CrcB library screens with a few exceptions. After washing, cells were inoculated into 200uL of both permissive and non-permissive media at a final dilution of 1/100. Three replicates of each ortholog and design candidate control were assayed. Microplate assays for each library were conducted by first randomly picking colonies following library transformation into design candidate gene knockout strains. After washing, one replicate of each colony and appropriate controls were inoculate into non-permissive growth media at a final dilution of 1/100. All microplate assays were conducted in 96 well plates. OD_600_ were measured in a BioTek Synergy HTX microplate reader with plates incubated with shaking at 30°C.

#### In vitro analysis of protein variants

To quantify flavinylation of SdhA by SdhE variants *in vitro*, SdhA, SdhE WT and SdhE variants were fused to a N-terminal 6xHis epitope tag, cloned into a TWIST-pT7-Kan inducible vector with a T7 RBS, expressed in *E. coli* BL21. To prevent *in vivo* flavinylation of SdhA, expression of SdhA was conducted in an *E. coli* BL21 *ΔsdhE* strain generated through P1 transduction. SdhE WT and SdhE variants were purified via Ni^+^ affinity chromatography. Total SdhE protein concentrations were calculated using ThermoFisher Quibit Protein Assay, and protein concentrations were normalized. Lysate from SdhA expressing cells was incubated with an equal amount of purified SdhE WT, a known non-functional SdhE point mutant, or SdhE variants in reactions with or without added flavin adenine dinucleotide (FAD). Reactions were visualized on SDS-PAGE using UV fluorescence to detect FAD and Coomassie stain to detect protein as previously described^25^. The co-localization of FAD with SdhA on SDS-PAGE gels confirmed *in vitro* covalent modification of SdhA by SdhE variants.

A fluoride riboswitch driven reporter system that provides heterologous validation of CrcB variant function was adapted from a previously described protocol^46^. A *Pseudomonas syringae* fluoride riboswitch driving the LacZ reporter was expressed in *E. coli* W3110 *ΔsdhE* strains complemented with CrcB variants. Overnight cultures of strains containing the riboswitch were grown overnight in LB supplemented with 2% glucose to suppress native LacZ, appropriate antibiotics, 10nM HSL inducer to express CrcB variants, and either 0.1mM or no NaF. In the morning, optical density of cultures was measured, cells were pelleted, and lysed in a modified Z-buffer (100 mM NaPO₄ pH 7.0, 10 mM KCl, 1 mM MgSO₄, 10% B-PER, 0.1 mg/mL lysozyme, 50 mM β-mercaptoethanol). Cell lysates were centrifuged and supernatant was transferred to a fresh plate. O-Nitrophenyl-β-D-galactopyranoside (OPNG) was added to each lysate to a final concentration of 1mg/mL. Reactions were incubated at 28°C in a plate reader, and OD₄₂₀, and Miller units were calculated using the formula (1000*OD_420_) / (T * V * OD_600_ * (1/D)) where T is time since adding OPNG, V is the volume of culture, and D is the culture dilution factor.

The enzyme kinetics of PanD variants was quantified through *in vitro* enzyme assays. PanD variants fused to 6xHis epitope tag in pBLxG2 vectors were expressed for 20 hours at 30°C in *E. coli* W3110 *ΔpanD* or *ΔpanDΔpanZ* and purified via Ni^+^ affinity chromatography. Since processing of PanD proenzyme is required for function^47^, and it is not feasible to separate processed PanD from proenzyme, processing of PanD was quantified by separating purified protein on SDS-PAGE and using ImageJ to estimate the proportion of processed enzyme. Total protein concentrations were calculated using ThermoFisher Quibit Protein Assay, and protein concentrations were normalized for *in vitro* enzyme assays. *In vitro* enzyme assays were conducted at 30°C using a variety of substrate concentrations. Reactions were quenched by addition directly to a Marfey’s regent reaction in acetone. Enzyme activity was determined using HPLC with detection at 340nm to quantify conversion of aspartate into β-alanine, and enzyme kinetics accounted were adjusted to the calculated concentration of processed PanD.

### Data analysis

#### Sequencing analysis

Adaptors were trimmed from paired end reads using Cutadapt^48^ and reads were aligned to reference files containing nucleotide sequences for all library designs using Bowtie2^49^. Total read counts were calculated for each protein variant in the library and converted to read count frequencies. For each biological replicates, the log2 enrichment scores were calculated by taking the log2 of read count frequencies for each variant in non-permissive growth media divided by read count frequences for each variant in permissive growth media. Log2 enrichments for each replicate were normalized to the log2 enrichment of the WT *E. coli* control, and the average and standard deviation was calculated across the three biological replicates for each library. Binary function labels were assigned relative to normalized log2 enrichment based on microplate assays, such that no variants considered functional were identified as non-functional when assayed individually.

#### Protein Scoring Metrics

ESMfold^34^ was used to predict structures of all protein variants and TM-scores were calculated using USAlign^37^ for each variant relative to the same *E. coli* WT structures specified for CCMpred^45^ constrained Potts Models. Model likelihood probabilities were calculated from pretrained ESM-1v^33^ and ESM-IF^36^ models using PyTorch^50^ and custom Python scripts. ProGen2^32^ likelihoods were obtained by averaging the mean of N-C terminus and C-N terminus likelihoods for each protein variant calculated using the progen2-large model. ProteinMPNN^35^ likelihoods were calculated using the v_48_020 model and ESMfold predictions of protein variants. HMM^30^ bits scores were calculated for each library using the profile HMM for the redesign candidate (PF02537 for CrcB, PF03937 for SdhE, and PF02261 for PanD). To calculate the percent ID to the nearest ortholog, a BLAST^51^ database was constructed from protein family MSAs, and the percent identity of the top BLAST hit for each design variant was selected. Potts model energy scores were calculated using a custom script. Potts models were built from MSAs by estimating single site frequencies and pairwise residue statistics across aligned positions. Pairwise coupling terms were estimating using direct coupling analysis. Fitted parameters were used to calculate energies for all MSA members and designed sequences.

Using a custom R script, ROC curves were generated with the binary functional label as the response, and AUC was calculated using the pROC package^52,53^. For comparison of AUC values in unobserved sequence space, libraries were split by percent identity to the nearest ortholog (for redesigned sequences) or percent ID to the *E. coli* WT sequence for natural orthologs such that equal numbers of functional sequences existed in both halves. This resulted in splits of 71.1%, 64.1%, 73% ID to natural orthologs for SdhE, CrcB, and PanD redesigns respectively and 42.5%, 44.8% 47.4% ID to *E. coli* WT for SdhE, CrcB, and PanD orthologs respectively. To determine if functional variants were scored higher than closest non-functional variants by sequence distance, a custom R script was used to calculate the fraction of pairs for which a predictor was better for the functional variant compared to the non-functional variant. A higher value was considered better for all metrics except Potts model energies and ProteinMPNN^35^ likelihoods, where a lower value is considered favorable.

Protein sequences were embedded in ESM-1b^54^, ESM-2^34^ and ProGen2^32^ pretrained models using full length sequences and custom Python scripts. Sequence embeddings were obtained by averaging the hidden-state vector across all positions in the protein sequence. A custom R script was used to compute pairwise distances between protein sequences using cosine distance in embedding space, and nearest neighbor was defined as the sequence with the smallest cosine distance. Nearest neighbor accuracy was determined by the fraction of sequences with nearest neighbors of the same functional class.

#### Identifying functional designs between orthologs

A custom R script was used to identify functional designs that lie between orthologs in sequence space. Pairwise Hamming distance matrices were calculated for all designs to all orthologs, and all orthologs to all orthologs. For each functional design, sequences “triplets” were retained containing two orthologs and one design where a design was closer to two ortholog than the two orthologs were to each other. Triplets for which the design to ortholog distances exceeded the ortholog-to-ortholog distance (path excess) by a Hamming distance of at most 50 for CrcB and PanD and 25 for SdhE were not considered between orthologs. Since multiple triplets often existed for each functional design, the highest scoring triplet was selected based on the lowest value for path excess added to the absolute value of the difference between the distance of each ortholog and the design (centeredness) weighted by a factor of 0.5. This prioritized designs that were genuinely between orthologs in sequence space, as close to that ortholog pairs cumulative distance overall, and centered between the two orthologs.

#### Sequence UMAP and protein fitness landscapes

To construct UMAP representations of sequence space, a MSA containing all natural orthologs and designed sequences was one-hot encoded for each protein family using a custom Python script. Gap characters were ignored during encoding to prevent discrimination between orthologs and designs based on alignment gaps. A UMAP embedding was computed for all sequences, and 2D UMAP plots were generated using a custom Python script. To construct protein fitness landscapes and prediction landscapes, UMAP coordinates from 2D plots were overlayed with normalized fitness, ProGen2 log likelihood, or Potts model energy values using a custom R script. To normalize fitness values between ortholog and design datasets, normalized log2 enrichments were scaled separately between -1 and 1, with 0 matching the threshold for function. A final scaled fitness value was created by combing scaled ortholog and design fitness values and averaging positive and negative control fitness values found in both datasets. ProGen2 and Potts model energy scores were aligned to the scaled fitness value using a simple linear model, and the predicted ProGen2 and Potts model energy scores when scaled fitness was equal to 0, were used as the reference point for predicted functional threshold. To generate 3D landscape surfaces, metrics were interpolated onto UMAP plots using the Akima^55^ package and smoothed with Fields package^56^. A mask was applied to fade the surface outside of regions in UMAP space lacking data.

## Data availability

The authors declare that the data supporting the findings of this study are available within the paper and its supplementary information files.

## Code availability

Custom code used to analyze data and generate figures will be made available following publication in peer reviewed journal.

## Acknowledgements

We would like to thank Dante Ricci for his assistance during assay development and data analysis and interpretation. This work was performed under the auspices of the U.S. Department of Energy by Lawrence Livermore National Laboratory under Contract DEAC52-07NA27344 (LLNL-JRNL-2021579)

## Author contributions

**ND**: Conceptualization, Data curation, Formal analysis, Investigation, Methodology, Visualization, Writing – original draft. **TD**: Conceptualization, Data curation, Funding acquisition, Investigation, Software, Resources. **RH**: Data curation, Software, Resources. **MF**: Investigation. **DJW**: Methodology, Investigation. **CKY**: Methodology, Software, Investigation. **DMP**: Conceptualization, Funding acquisition. **SPL**: Conceptualization, Funding acquisition, Project administration, Supervision, Writing – original draft. **All authors**: Writing – review & editing.

## Additional information

Supplementary Information is available for this paper. Correspondence and requests for materials should be addressed to SPL.

## Extended data figures and tables

**Extended Data Figure 1.**
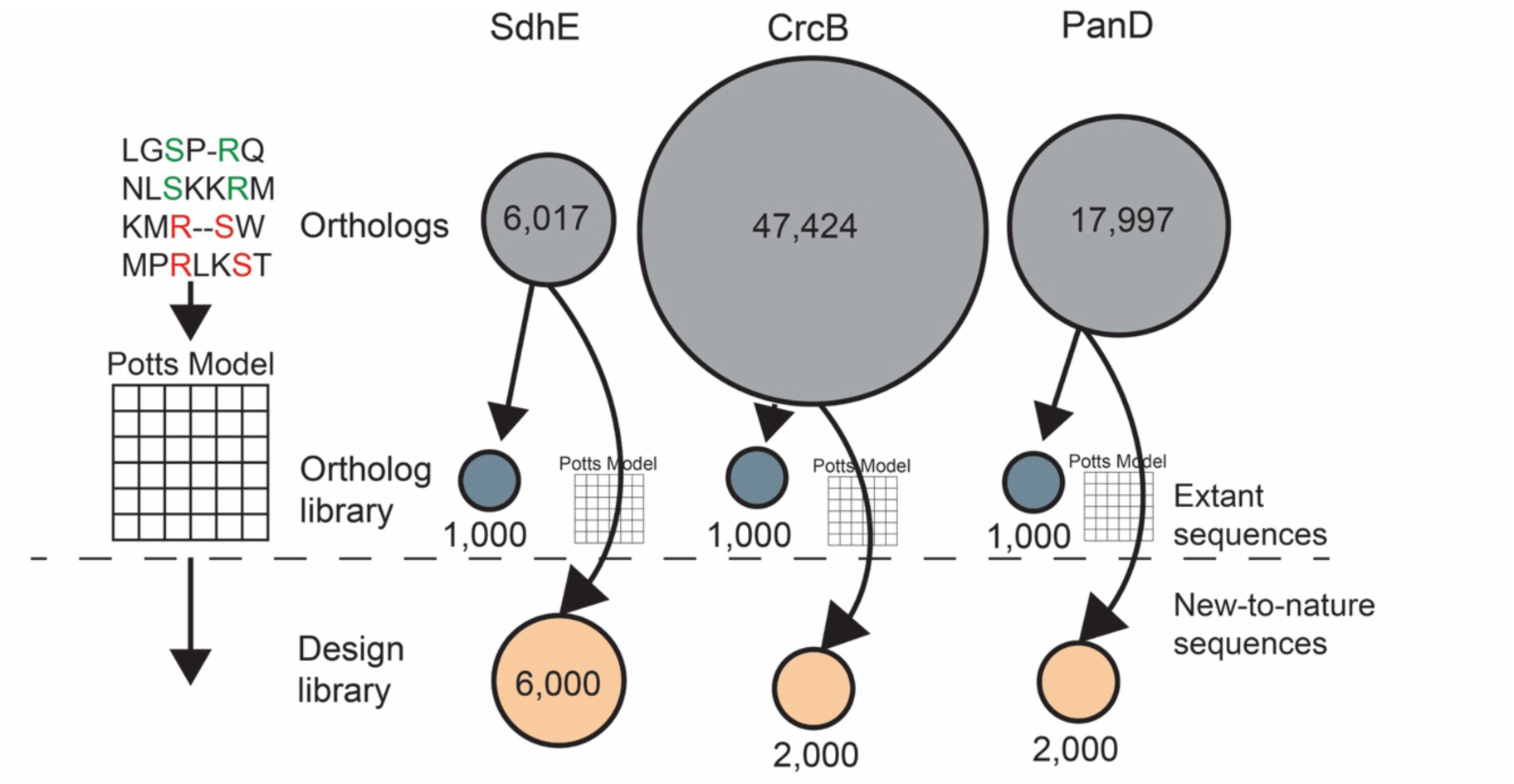
Overview of the size of observed sequence space relative to experimentally tested orthologs and new-to-nature Potts model designs.

**Extended Data Figure 2.**
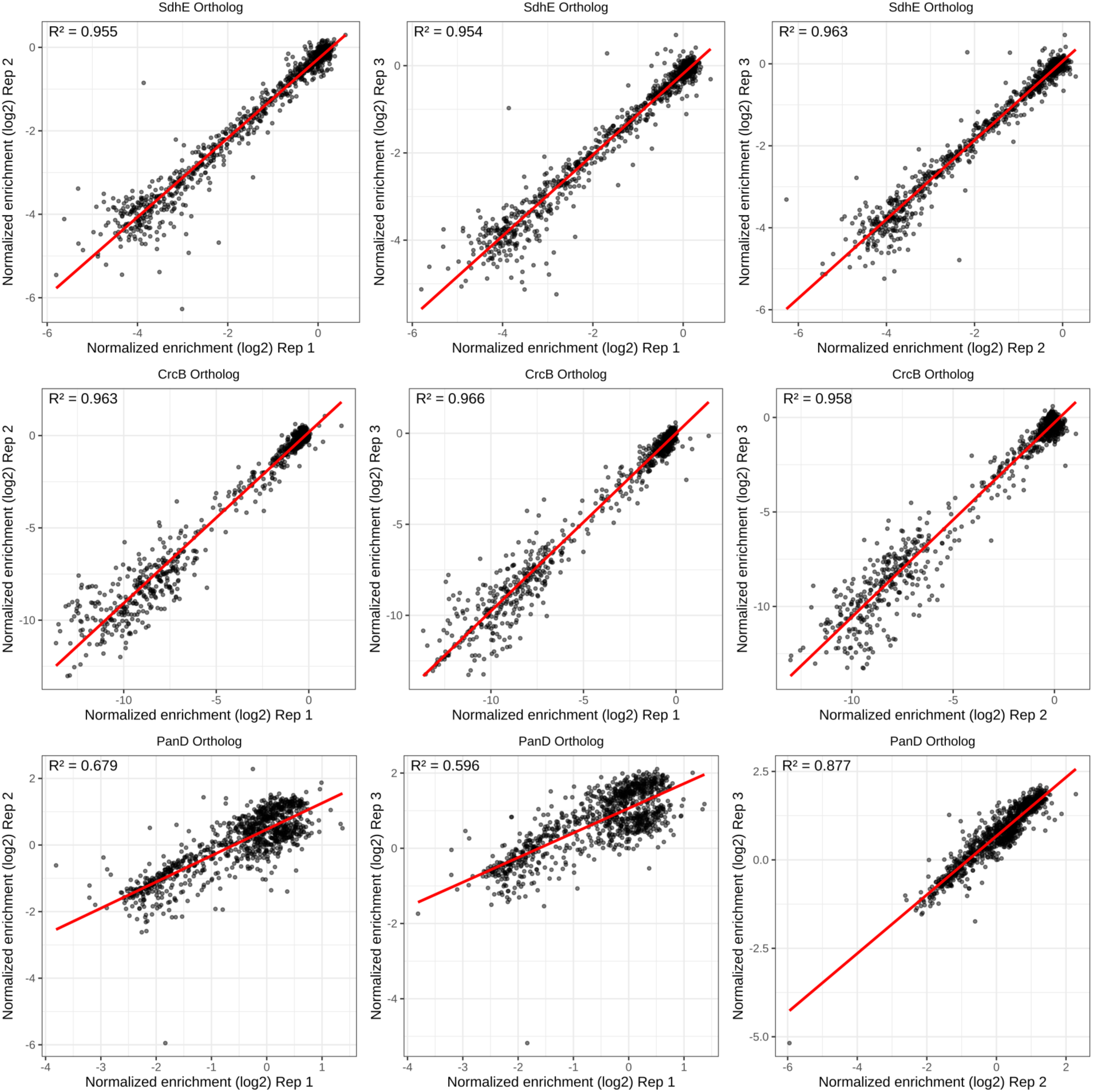
Correlations between normalized log2 enrichment of each of the three replicates for pooled growth-based screens for all ortholog libraries.

**Extended Data Figure 3.**
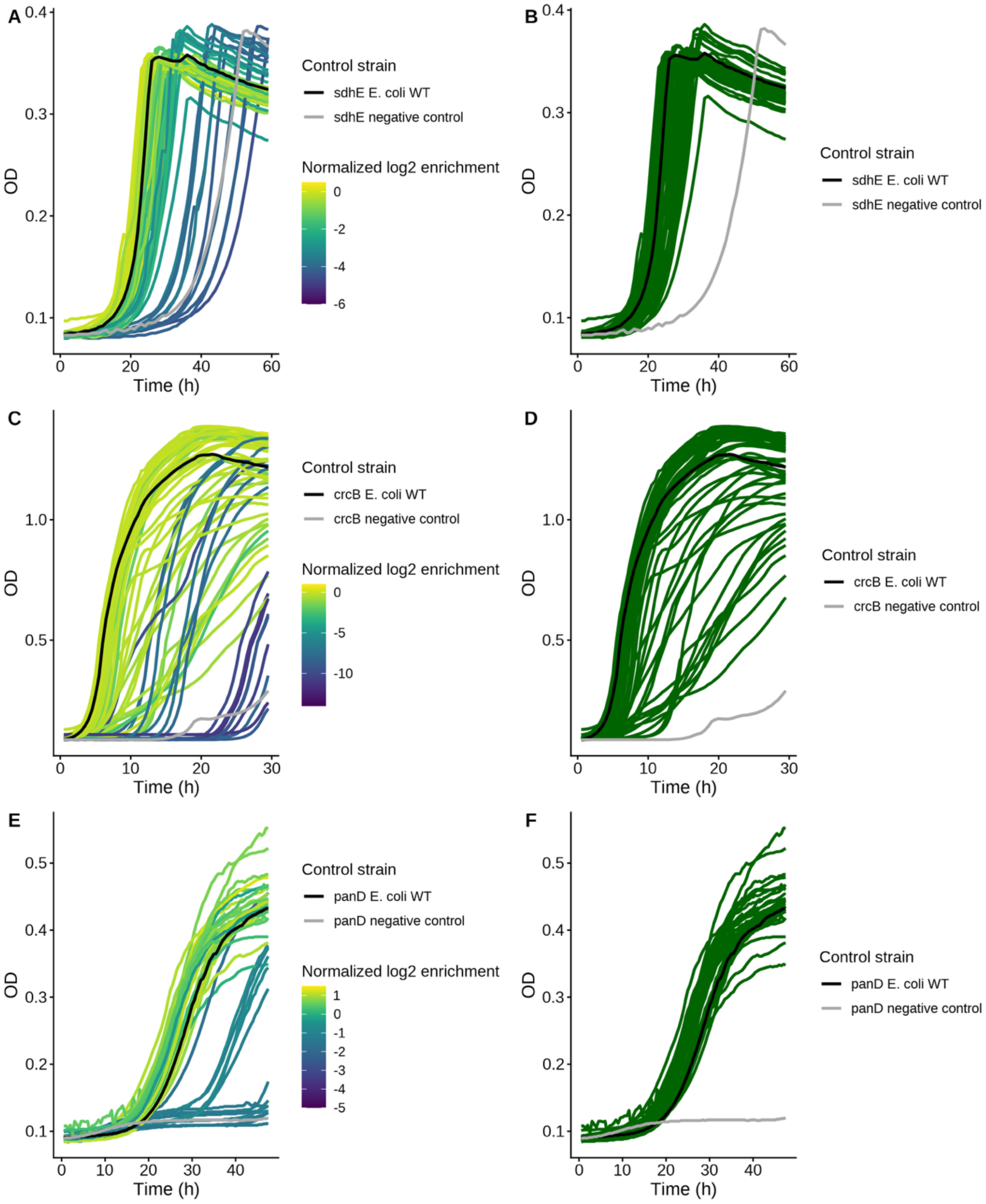
A subset of tested orthologs supports functional thresholds determined from pooled library assays when grown individually. A random selection of variants were grown in non-permissive media in microwell plates from SdhE (A, B), CrcB (C, D) and PanD (E, F) ortholog libraries (SdhE n=66, CrcB n=85, PanD n=44). A conservative normalized log2 enrichment functional threshold was determined such that functional variants (right column) were all able to grow in non-permissive media, while allowing for the possibility that certain variants determined to be non-functional based on library enrichment could function individually.

**Extended Data Figure 4.**
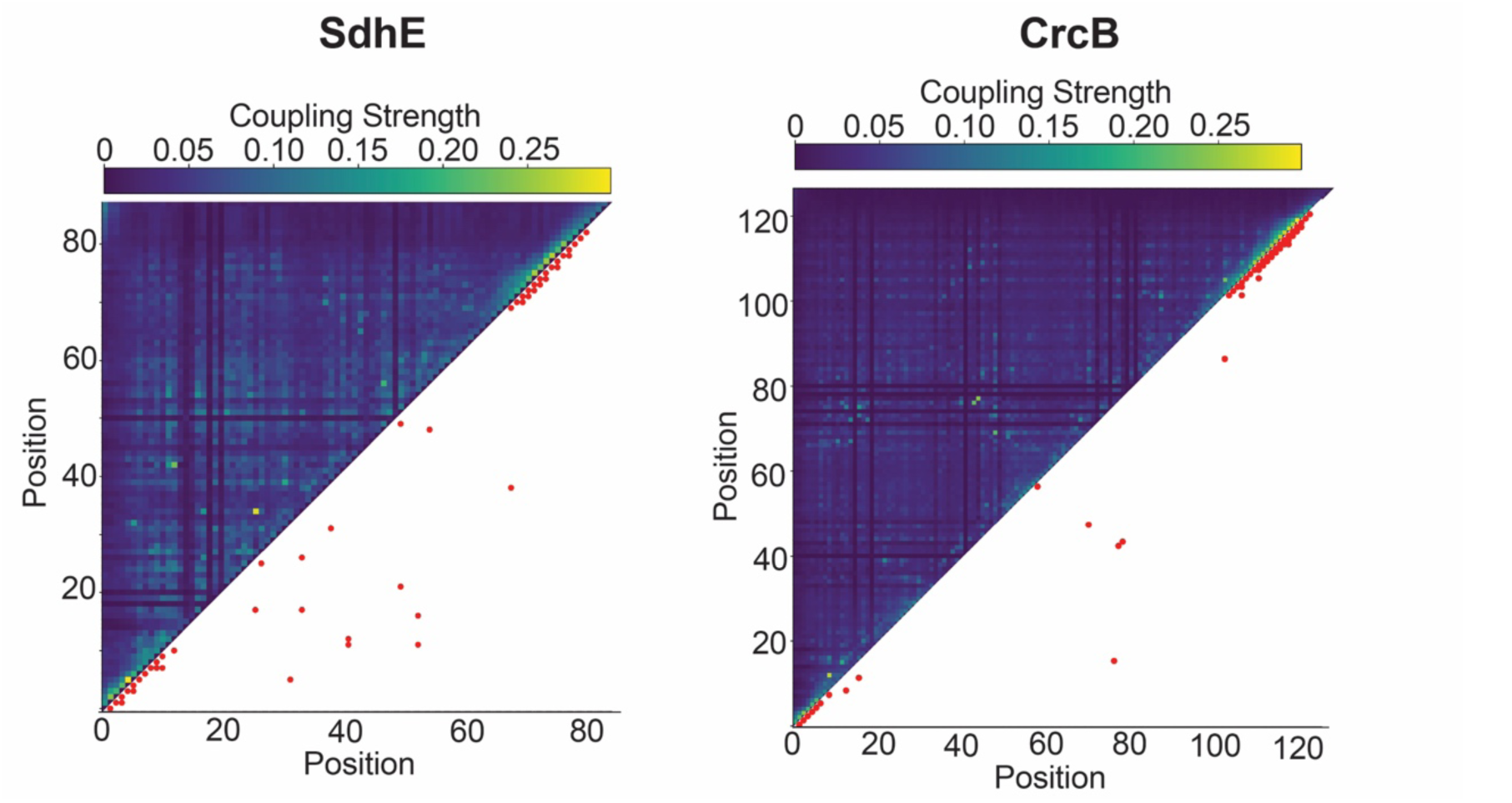
Coupling strength of all SdhE (left) and CrcB (right) positions based on direct coupling analysis with the top 50 strongest couplings represented by red points in the bottom right of each panel.

**Extended Data Figure 5.**
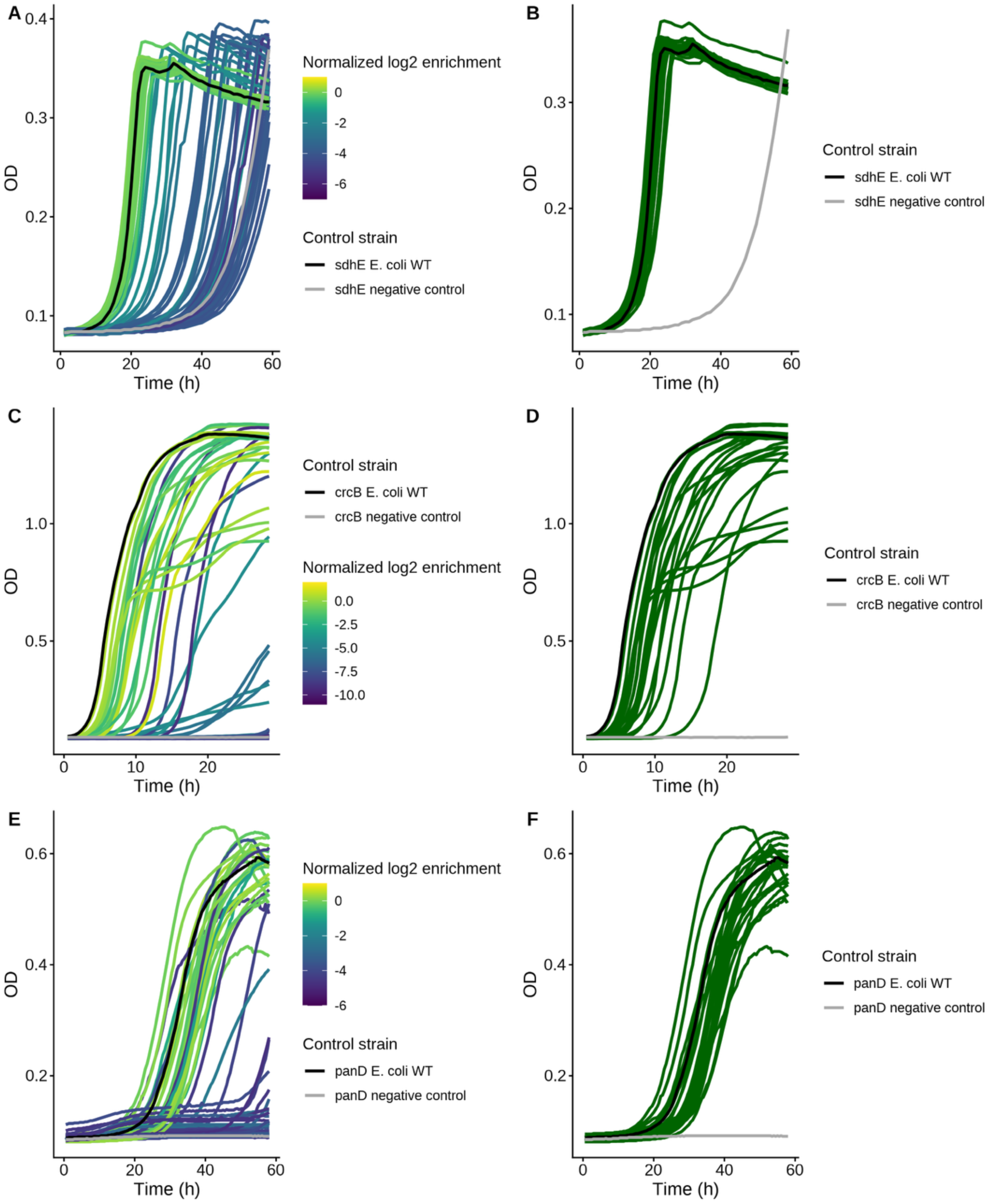
A subset of tested designs supports functional thresholds determined from pooled library assays when grown individually. A random selection of variants were grown in non-permissive media in microwell plates from SdhE (A, B), CrcB (C, D) and PanD (E, F) design libraries (SdhE n=68, CrcB n=71, PanD n=62). A conservative normalized log2 enrichment functional threshold was determined such that functional variants (right column) were all able to grow in non-permissive media, while allowing for the possibility that certain variants determined to be non-functional based on library enrichment could function individually.

**Extended Data Figure 6.**
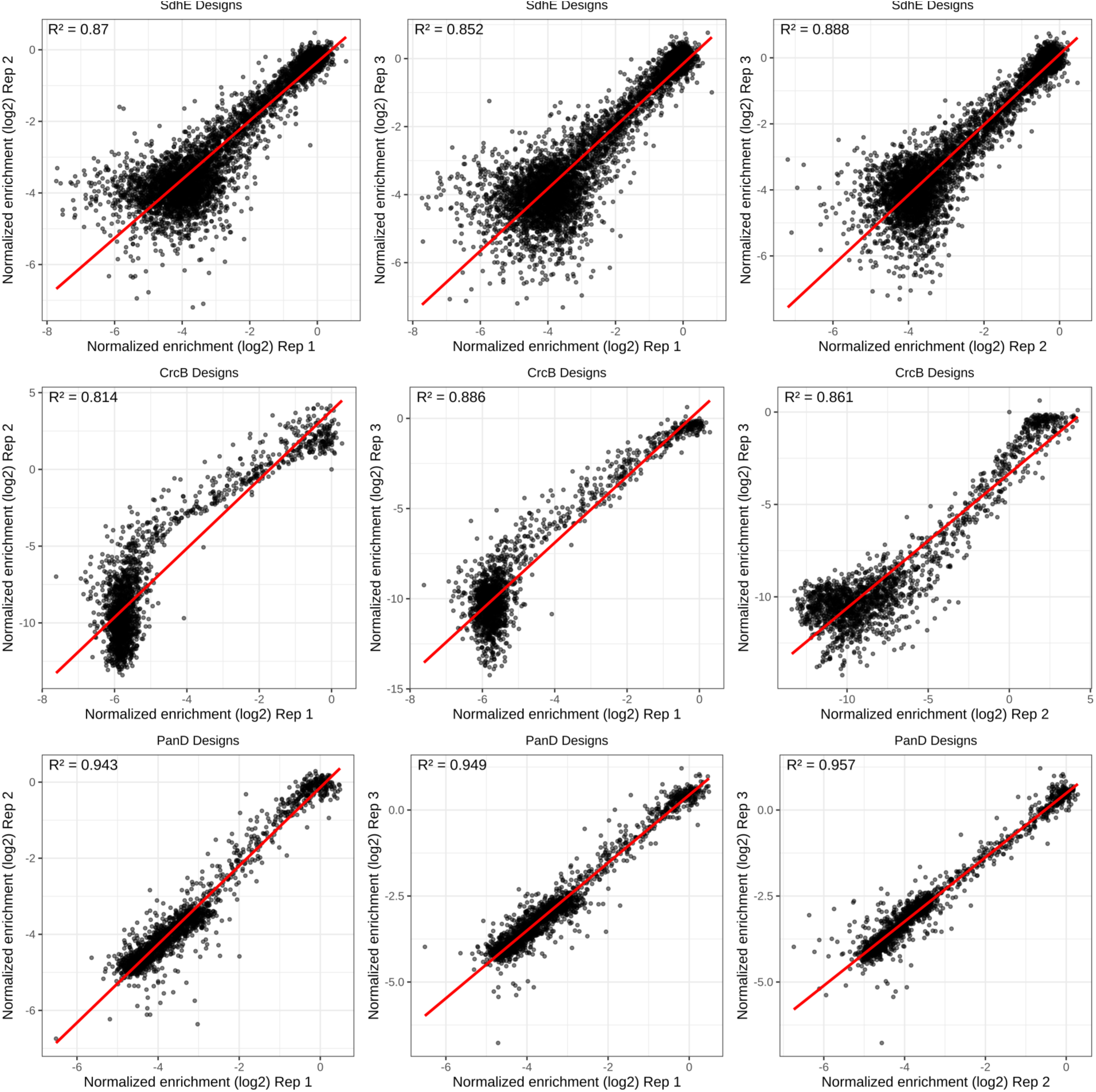
Correlations between normalized log2 enrichment of each of the three replicates for pooled growth-based screens for all design libraries.

**Extended Data Figure 7.**
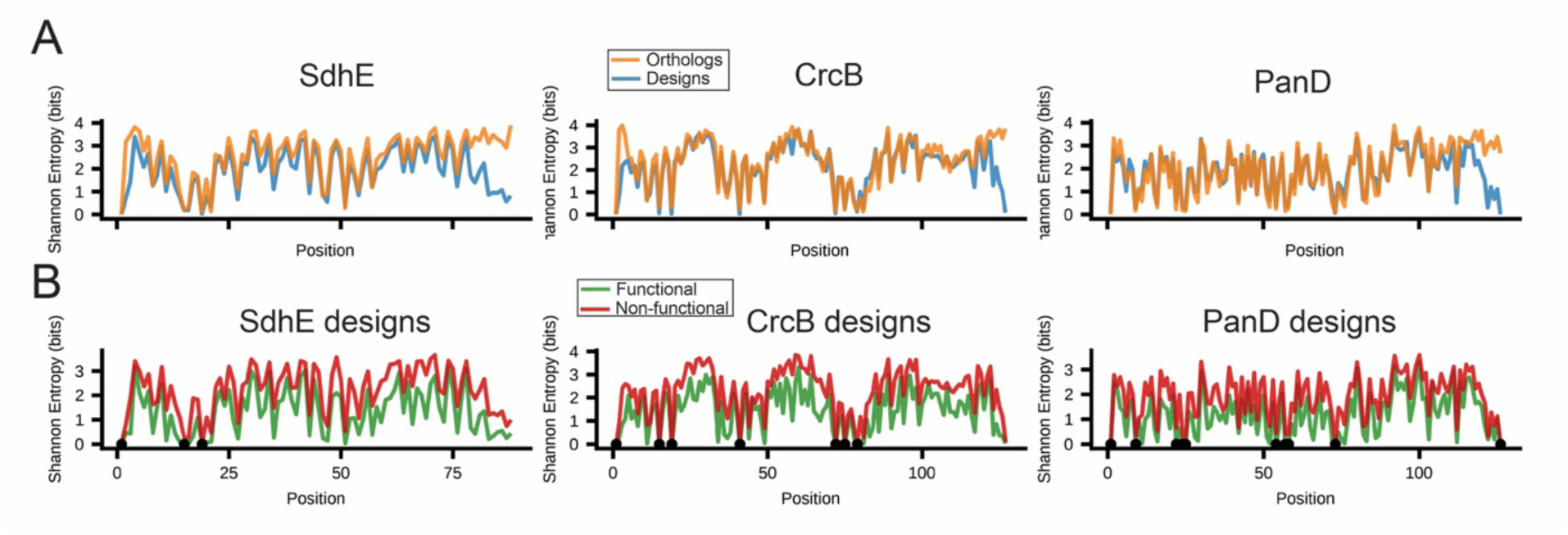
Shannon entropy of protein orthologs and designs. (A) Protein designs and orthologs have similar entropy across all positions across each of the three protein families. (B) Non-functional new-to-nature proteins generally have higher entropy than functional dewsigns, and all functional new-to-nature proteins are conserved at key conserved residues. Black dots on the x-axis indicate positions where entropy is 0 for all functional protein designs.

**Extended Data Figure 8.**
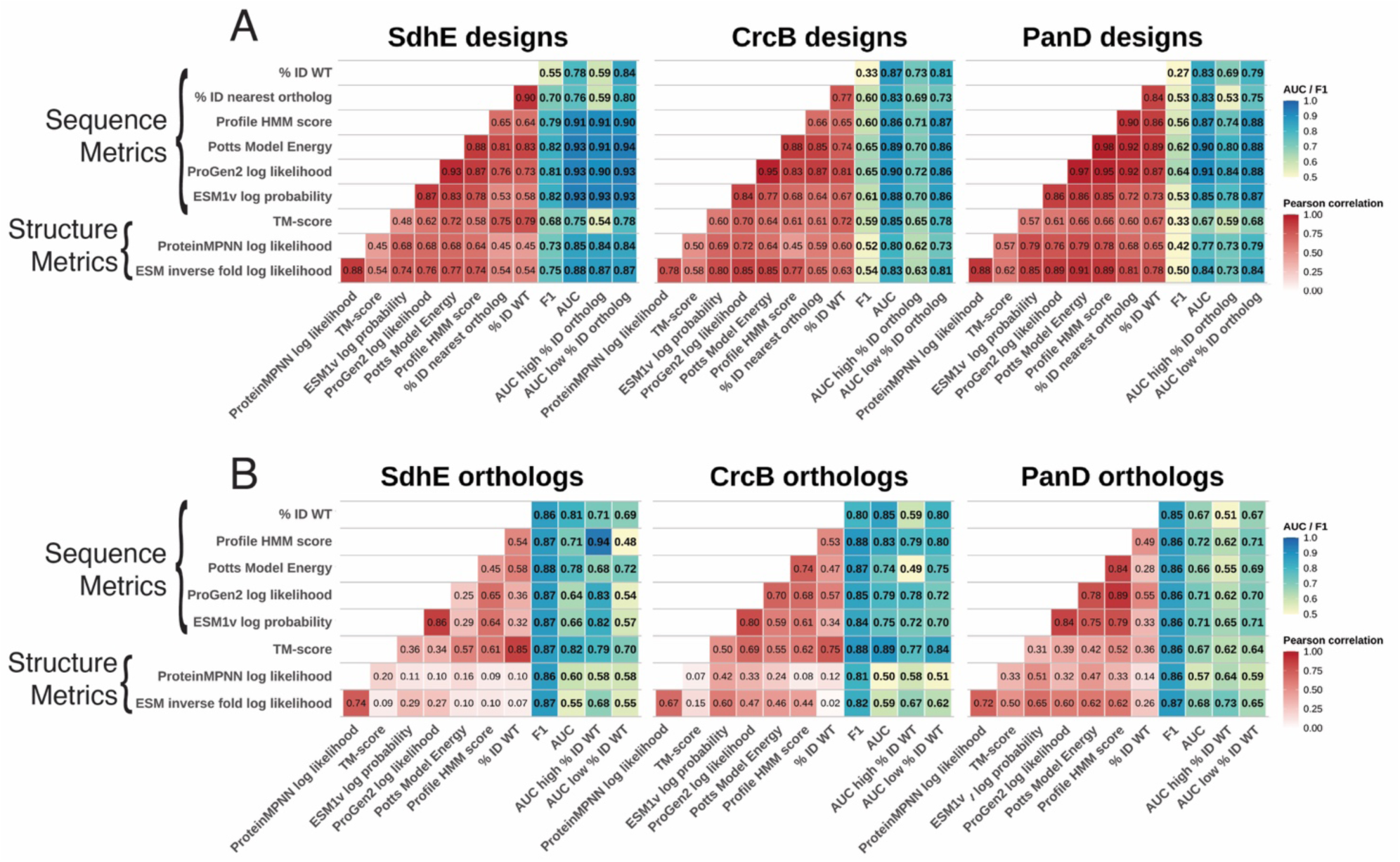
Protein scoring metrics predict function across the six tested libraries with varying accuracy. Diverse sequence and structure metrics pairwise absolute Pearson correlation, and performance on when function is represented as a binary determined via F1 scores, and AUC for (A) designs and (B) orthologs.

**Extended Data Figure 9.**
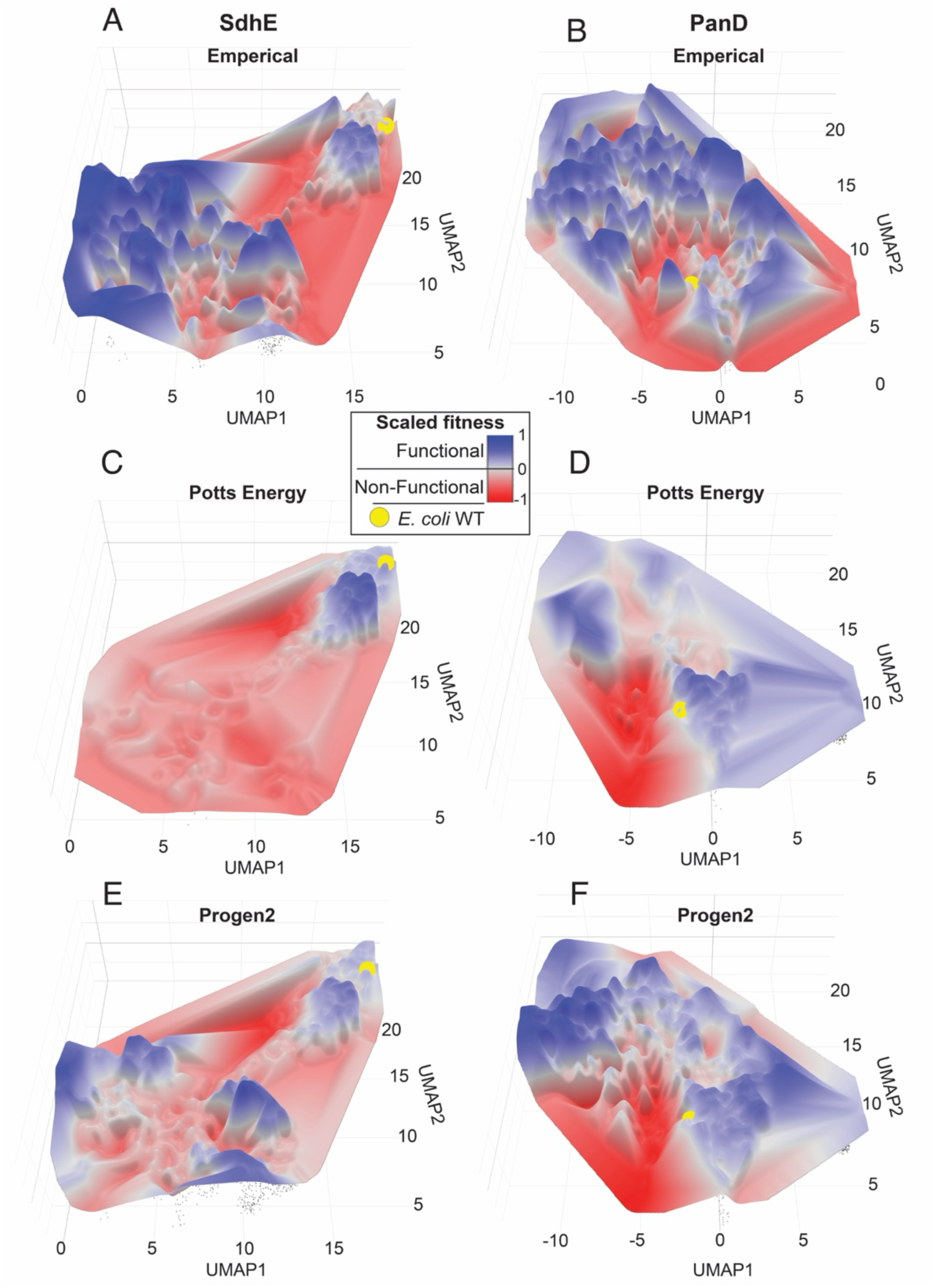
The SdhE and PanD protein fitness landscapes are rugged. UMAP representations of one-hot encoded SdhE (left column; A,C,E) and panD (right column; B,D,F) designs and natural orthologs on the x- and y-axis are overlayed with scaled fitness values from ortholog and design pooled assays (A,B). Scaled scores from Potts model energy (C,D) and ProGen2 (E,F) for all tested SdhE and PanD orthologs and designs are overlayed on the z-axis of the sequence UMAP plots. The *E. coli* WT protein sequence is shown as a yellow sphere. Fitness is scaled such that 0 (white) is the threshold for function across empirical and predicted landscapes.

**Extended Data Table 1.** Ruggedness metrics across fitness landscapes.

| Gene | Landscape | Roughness | Mean Gradient |
| --- | --- | --- | --- |
| SdhE | **Experimental** | **0.48** | **0.47** |
|  | ProGen2 | 0.32 | 0.30 |
|  | Potts Energy | 0.32 | 0.17 |
| CrcB | **Experimental** | **0.60** | **0.62** |
|  | ProGen2 | 0.25 | 0.31 |
|  | Potts Energy | 0.18 | 0.31 |
| PanD | **Experimental** | **0.44** | **0.55** |
|  | ProGen2 | 0.27 | 0.32 |
|  | Potts Energy | 0.26 | 0.21 |

