## Supplemental Information for "Unobserved Sequence Space Has Many Functional Proteins"

*In vitro and heterologous validation confirm function of unobserved proteins*

To validate that growth-based complementation of our tested proteins was due to correct protein function, we conducted additional *in vitro* validation assays on a subset of designed variants. To validate that SdhE variants are functional in promoting flavinylation of SdhA, we conducted *in vitro* flavinylation assays using purified SdhE variants, and cell lysate from *E. coli ΔsdhE* overexpressing SdhA as previously described[25]. We selected variants that were distant from and similar to *E. coli* SdhE WT based on sequence identity. We verified that functional variants could promote flavinylation of SdhA *in vitro* (Supplemental Fig. 1).

To determine if CrcB designs that complement growth in *E. coli ΔcrcB* are function fluoride efflux channels, we leveraged a previously described LacZ $F^-$ riboswitch assay[26,46]. We calculated Miller Units from cell lysates of *E. coli ΔcrcB* expressing the $F^-$ riboswitch LacZ construct, complemented with each of the tested CrcB variants used to determine functional thresholds in our CrcB design library pooled assays (Extended Data Fig. 5). When strains were grown in media with fluoride, low Miller Units corresponding to low intracellular fluoride correlated with normalized log2 enrichment above the functional threshold for CrcB (Supplemental Fig. 2). This directly supports the robustness of our growth-based assay by correlating log2 enrichment with protein function.

To validate that PanD variants function as aspartate decarboxylases, we conducted *in vitro* enzyme assays using purified PanD variants. Our selected variants included 8 functional and 1 non-functional variant based on our library screen assay. We grew *E. coli ΔpanD* complemented with variants in non-permissive media to validate that function of each variant was not affected by an N-terminal 6x-His tag and found that 7/8 functional variants and the 1 selected non-functional variant could complement growth in non-permissive conditions (Supplemental Fig. 3). PanD is expressed as a zymogen that must undergo processing to form a functional enzyme comprised of a small (α) and large (β) subunit[47]. While this processing is promoted by PanZ in *E. coli*, and *E. coli ΔpanZ* replicates the conditionally essential growth phenotype of *E. coli ΔpanD*[50], not all organism require a PanZ ortholog for processing[47]. In fact several PanD orthologs have been described which undergo processing efficiently in the absence of PanZ[51].

We found that all tested PanD variants could catalyze the formation of β-alanine from L-aspartate, but that the catalytically dead $PanD^{S25A}$ mutant, which is also not capable of processing into small and large subunits, could not form detectable β-alanine. We also found that enzyme parameters did not correlate well with normalized log2 enrichment (Supplemental Fig. 4). This lack of correlation is likely due to the proportion of enzyme processing being factored into our calculation of enzyme kinetics (Supplemental Fig. 4) which may misrepresent the amount of β-alanine being produced *in vivo* in the context of our growth-based assays. To explore pro-enzyme processing further, we expressed PanD variants in *E. coli ΔpanDΔpanZ*. While processing was significantly reduced for PanD WT in the absence of PanZ, several variants were able to process in the absence of PanZ (Supplemental Fig. 4). Although examples of natural PanD orthologs that can process without PanZ exist[51], the presence of auto-processing in our PanD designs demonstrates that protein redesign can create new functions relative to the *E. coli* PanD WT starting point. Only one of the evaluated panD designs had greater kcat than the *E. coli* PanD WT, suggesting opportunities for further improvement towards enzyme redesigns with greater catalytic activity.

### Function prediction of orthologs and designs

Our finding that Potts model energy scores match or outperform pLM log likelihoods led us to investigate correlations between scoring metrics. Potts model energy scores are highly correlated with Progen2 log likelihoods (Extended Data Fig. 8, Supplemental Fig. 5). This suggests that either pLMs capture similar information to lightweight family specific Potts models, which has been suggested previously[52], or that both scoring metrics arrive at highly similar predictions of function using different sequence-based information. We also found that ESM1v and Progen2 log likelihoods as well as ProteinMPNN and ESM-IF likelihoods were highly correlated across the tested design libraries (Supplemental Fig. 6, Supplemental Fig. 7)

We wanted to further explore low accuracy of function prediction across ortholog libraries. Given that natural ortholog sequences are present in all evaluated model training data, this is likely due to favorable scores being assigned across a large proportion of orthologs. We observe higher average scores for orthologs relative to new-to-nature proteins for all tested AI metrics and Potts model energies, but not HMM scores (Supplemental Fig. 8).

To determine if AI-based protein metrics could generalize into novel unobserved sequence space, we evaluated each metric on our designed proteins when split by percent ID to the nearest ortholog such that both halves have equal numbers of functional sequences. Scoring sequences using a protein family profile HMM, Potts model, Progen2 or ESM1v resulted in similar AUC values for proteins with low identity to extant sequences within all three protein families (Extended Data Fig. 8). Metrics generally had better accuracy on designs with low identity to natural orthologs, although this is likely due to class imbalance (Extended Data Fig. 8). No metrics consistently predicted function of orthologs with low similarity to the *E. coli* WT sequence (Extended Data Fig. 8) Overall, our results suggest that AI protein tools do not generalize into unobserved sequence space better than simple lightweight scoring metrics (Extended Data Fig. 8).

Previous research suggests protein language model embeddings can group proteins with similar function despite low sequence similarity[53,54]. We extracted ESM1b, ESM2, Progen2 small-model, and Progen2 large-model embeddings for all sequences in our libraries and measured sequence distance or cosine distance in embedding space. New-to-nature proteins for all three genes had a higher proportion of nearest neighbors in embedding space of the same functional class when compared to sequences space, suggesting that embeddings capture more information about protein function than sequence distance alone (Supplemental Fig. 9). ESM2 generally matched or slightly exceeded the performance of ESM1b and both ESM models exceeded performance of both Progen2 models (Supplemental Fig. 9). While Progen2 embeddings cluster nearest neighbors by function in embedding space slightly better than sequence distance alone for new-to-nature libraries, sequence distance outperforms Progen2 models for SdhE and CrcB ortholog libraries (Supplemental Fig. 9). Moreover, distance in UMAP space using one-hot sequence encodings clusters sequences by functional class more accurately than Progen2 models for half of the tested libraries (Supplemental Fig. 9). This suggests that simple embeddings and dimension reduction can recover much of the accuracy captured by pretrained Progen2 models, but that nearest sequences in ESM embedding space are more likely to have identical function when compared to UMAP sequence space or sequence distance.

### Distribution of function in unobserved sequence space

We set thresholds to determine which designs were between orthologs in sequence space based on hamming distances and accounting for the centeredness and total excess distance of paths between two designs through a given ortholog (see methods). Being centered is important in this context, since a perfectly centered new-to-nature protein indicates that each variable positions in the triplet is different in only one ortholog, and common between the new-to-nature protein and other ortholog (Figure 4A). Therefore, in perfectly centered triplets, a set of mutations could be introduced in one ortholog to reach a new-to-nature protein, and a completely independent set of mutations could be introduced into the new-to-nature protein to reach the remaining ortholog, indicating that new-to-nature proteins are literally between occupied sequence space. We found that 52%, 65%, and 28% of unobserved sequence space between occupied sequence space was functional for SdhE, CrcB, and PanD respectively (Supplemental Table 1).

The remaining functional protein designs across all three protein families are either near natural orthologs or on distant indirect paths outside of occupied sequence space. These designs can be defined as not satisfying the geometric constraints (the distance between the design and either ortholog must be less than the distance between either ortholog), path excess (the design to ortholog distances exceeded the ortholog-to-ortholog distance by a Hamming distance of at most 50 for CrcB and PanD and 25 for SdhE), or both. Exceeding the path excess means the design lies in distant sequence space, with trajectories that are far from straight paths between natural orthologs. To identify functional designs that were distant from natural orthologs in sequence space, we focused on functional designs that did not satisfy the path excess constraint, or both constraints. We identified 1326 functional SdhE designs, 91 functional CrcB designs, and 18 PanD designs that did not satisfy the path excess constraint or the path excess and geometric constraint.

### Supplemental Data

Supplemental data will be made available upon publication in a peer reviewed journal.

**Supplementary data S1**: Next-generation sequence read counts and coverage for all library samples

**Supplementary data S2**: Protein variants tested in plate reader assays across all ortholog and design libraries

**Supplementary data S3**: Tested proteins with normalized log2 enrichment scores and functional labels from pooled growth-based assays across all libraries

### Supplemental figures and tables

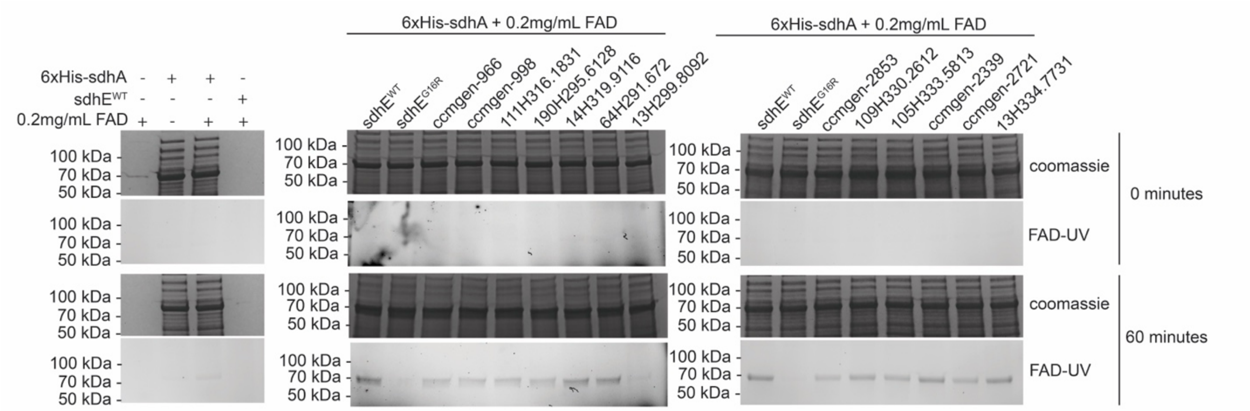


Supplemental Figure 1. *In vitro* validation of growth based functional complementation of select SdhE designs. Purified SdhE promotes flavinylation of SdhA from lysate of *E. coli BL21 ΔsdhE*. Initial and 1-hour timepoints of *in vitro* sdhA flavinylation assays were separated via SDS-PAGE. FAD was visualized using UV-light, and protein was visualized through Coomassie stain. Selected SdhE variant are distant from natural ortholog sequences (center) or similar to natural ortholog sequences (right).

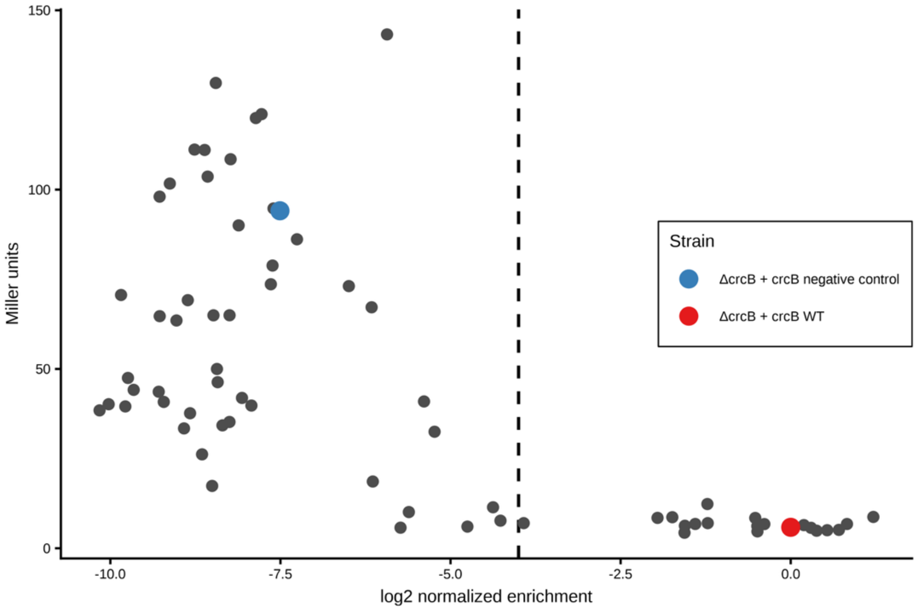


Supplemental Figure 2. Heterologous validation of growth based functional complementation of select CrcB designs. Miller assay of *E. coli ΔcrcB* complemented with a randomly selected pool of new-to-nature variants with fluoride riboswitch LacZ reporter constructs. Miller units increase below the functional threshold (vertical dotted line) indicating an increase in intracellular fluoride due to lack of CrcB complementation.

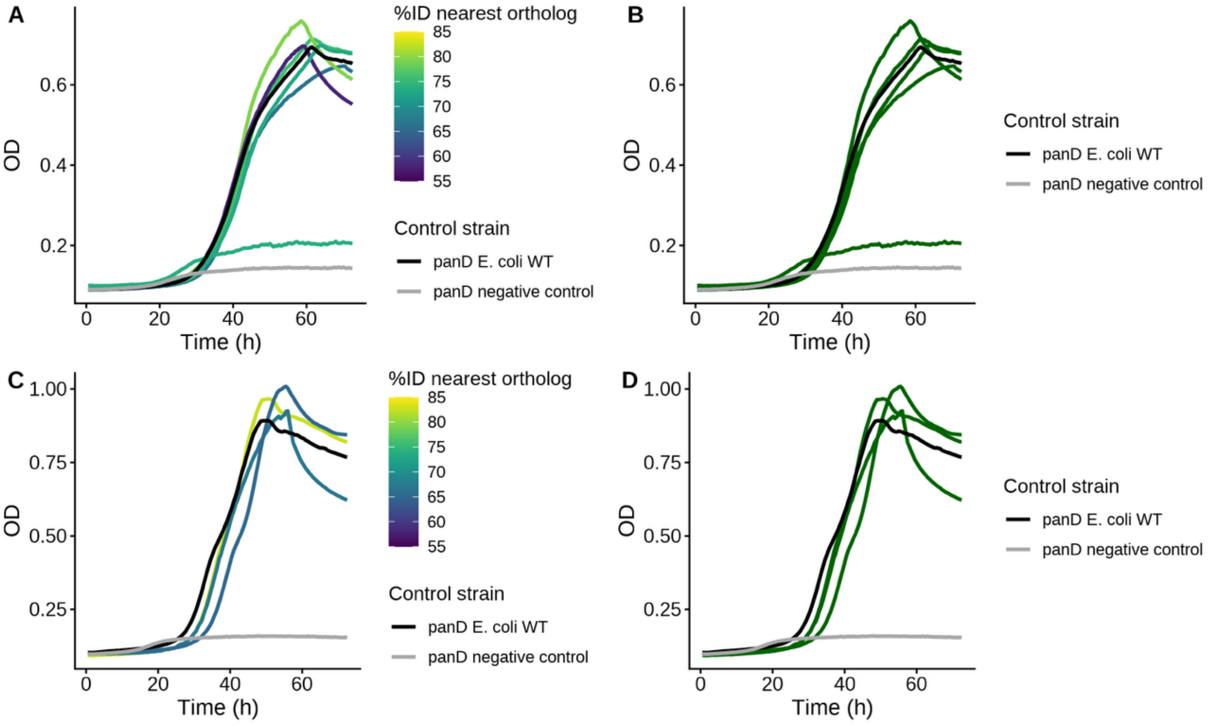


Supplemental Figure 3. Expression of panD variants with N-terminal 6x-His fusion used for *in vitro* enzyme assays complement *E. coli ΔpanD* in non-permissive media. Variants have diverse similarity to existing natural orthologs (A,C) and functional variants as determined by library based normalized log2 enrichment threshold (B,D).

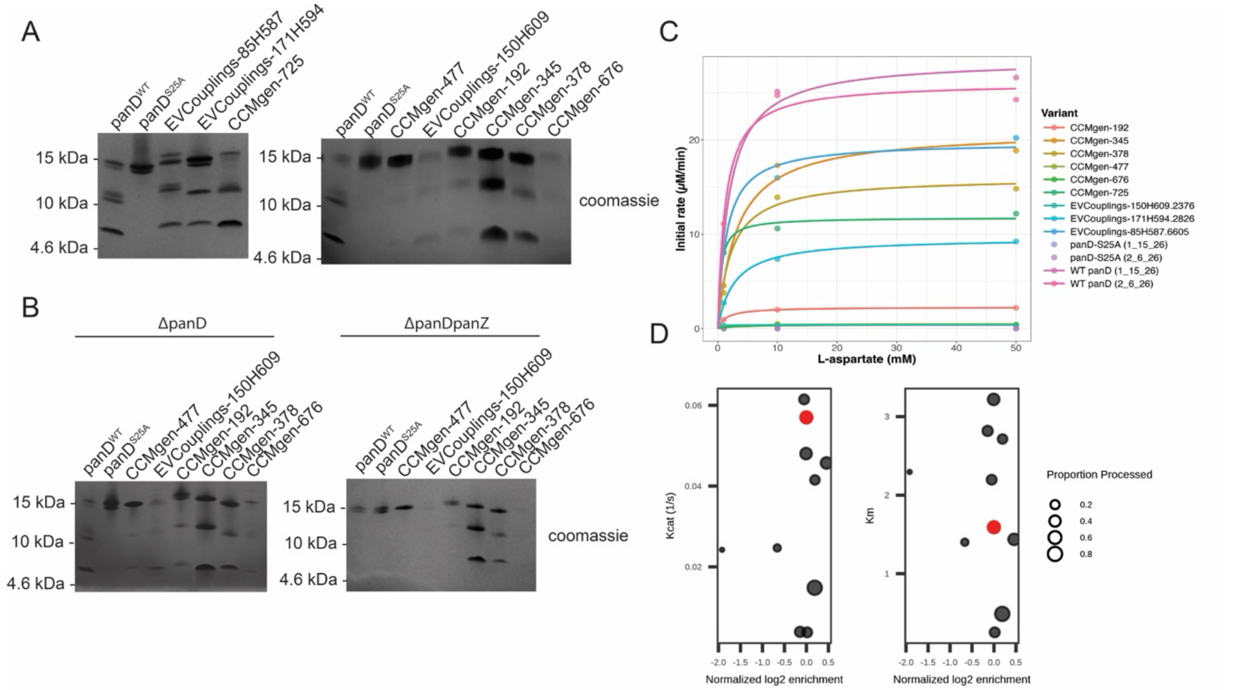


Supplemental Figure 4. In-vitro validation of growth based functional complementation of select PanD designs. Select PanD variants show differing abilities to process in (A, B left) *E. coli*

*ΔpanD,* and (B right) *E. coli ΔpanDΔpanZ*. PanD variants were fused to an N-terminal 6xHis tag and purified via affinity chromatography. Purified variants were separated via SDS-PAGE and visualized with Coomassie stain, and band intensity (A) was used to quantify the proportion of processed versus proenzyme PanD. (C) Michaelis-Menten kinetics curves for PanD variants assayed *in vitro* (D) k-cat and Km values determined for PanD variants through *in vitro* enzyme kinetic assays, based on normalized log2 enrichment. The size of dots indicates the amount of processed panD enzyme relative to total panD protein (A).

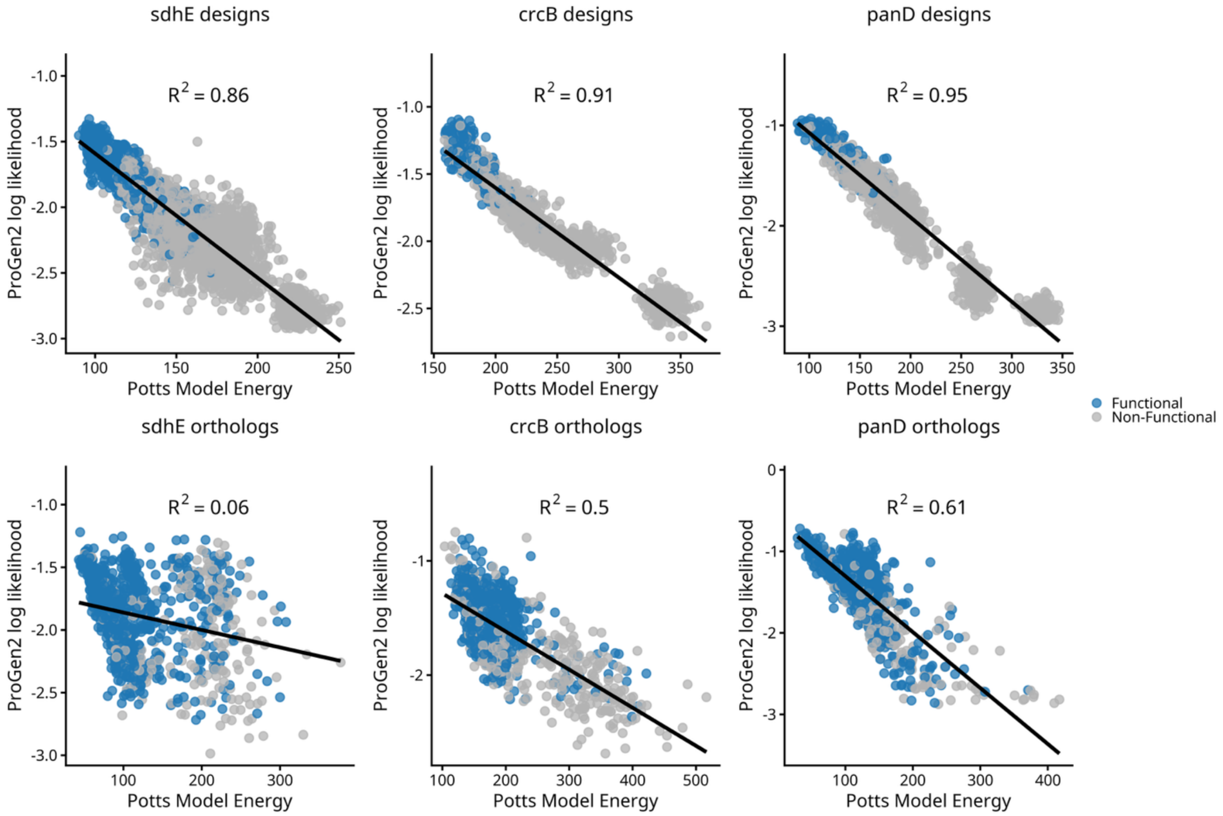


Supplemental Figure 5. Potts model energy scores are highly correlated with Progen2 log likelihood scores in design libraries (top rows) but not ortholog libraries (bottom row). Black line is linear regression trendline, blue dots indicate functional library variants while grey dots indicate non-functional variants.

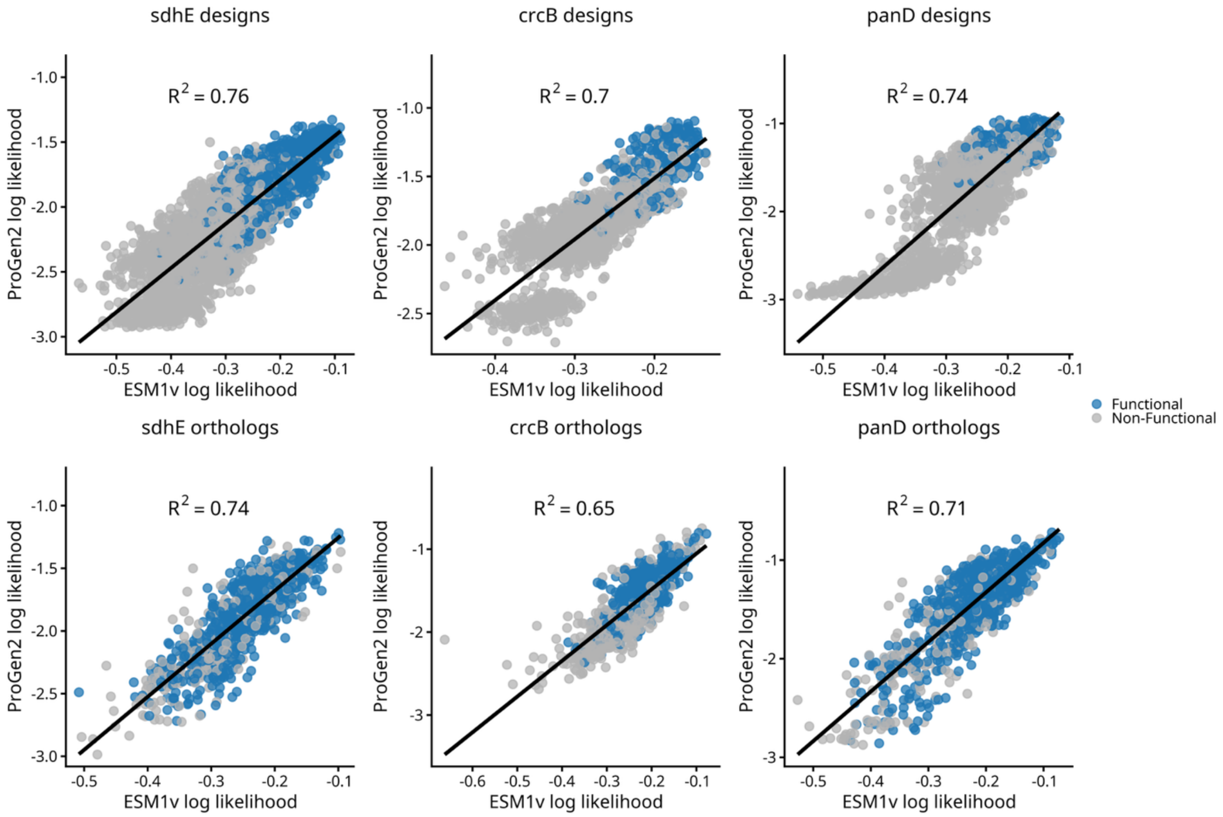


Supplemental Figure 6. Likelihoods from Progen2 and ESM1v protein language models are correlated across all tested design (top row) and ortholog (bottom row) libraries. Black line is linear regression trendline, blue dots indicate functional library variants while grey dots indicate non-functional variants.

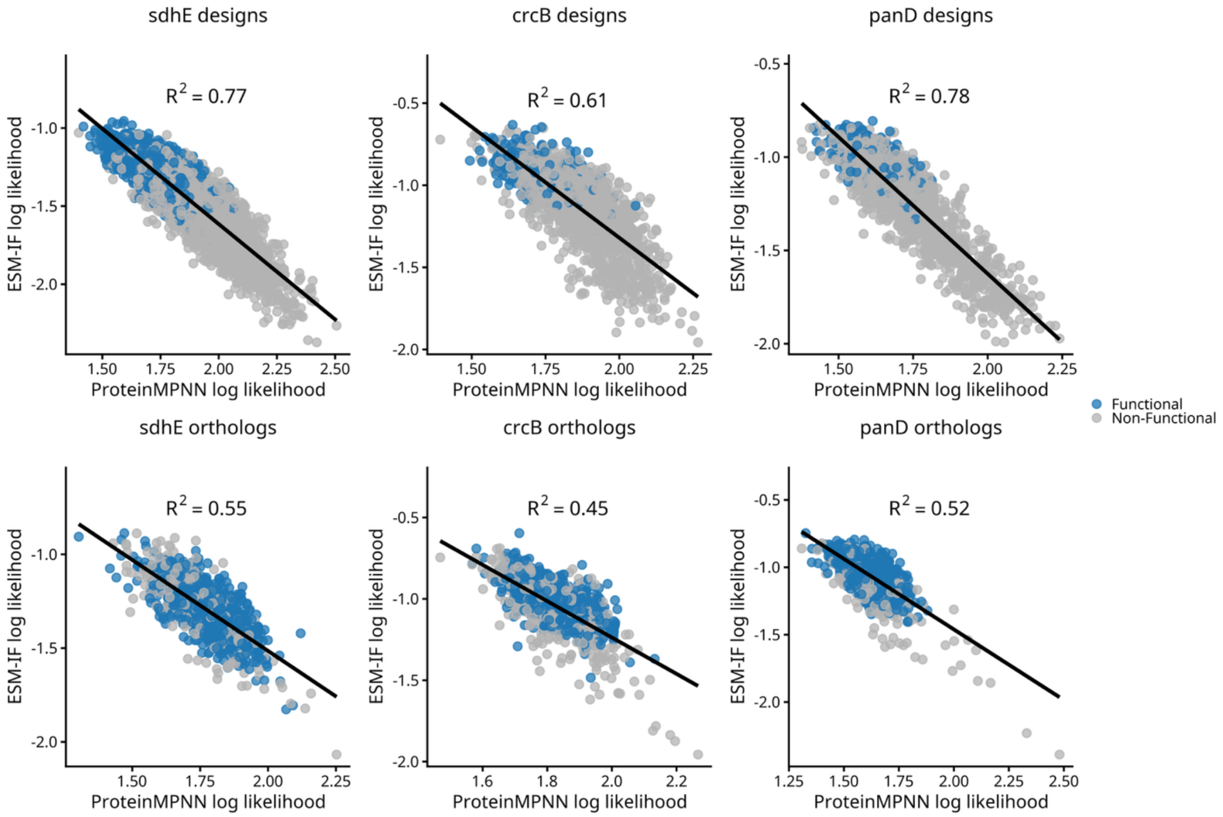


Supplemental Figure 7. Likelihoods from ProteinMPNN and ESM Inverse Fold (IF) models are correlated across all tested design (top row) and ortholog (bottom row) libraries. Black line is linear regression trendline, blue dots indicate functional library variants while grey dots indicate non-functional variants.

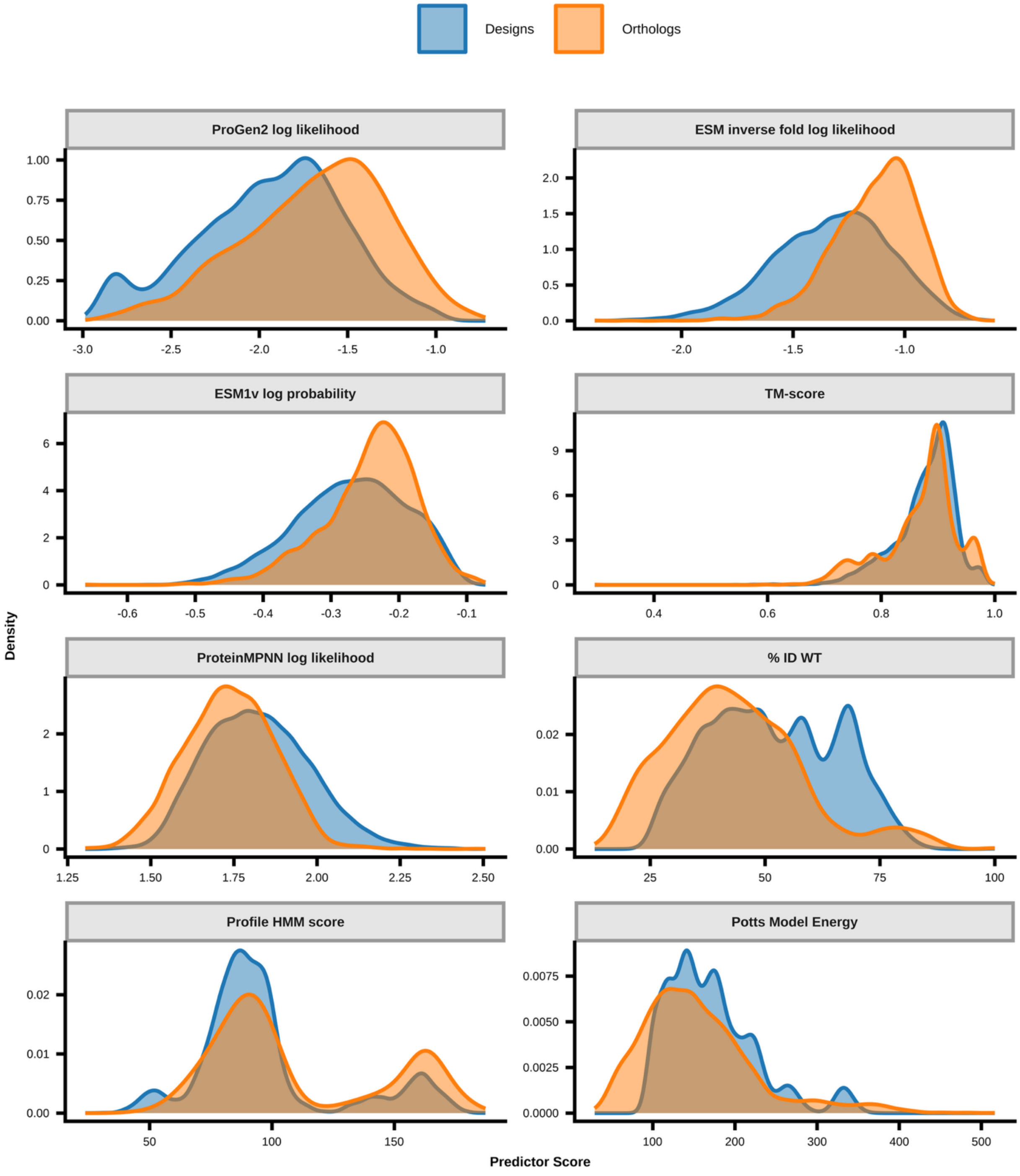


Supplemental Figure 8. Distribution of scores across several metrics for combined SdhE, CrcB, and PanD ortholog and design libraries. Orthologs have greater densities of higher scores relative to designs for AI-based metrics (top two rows) relative to non-AI-based metrics.

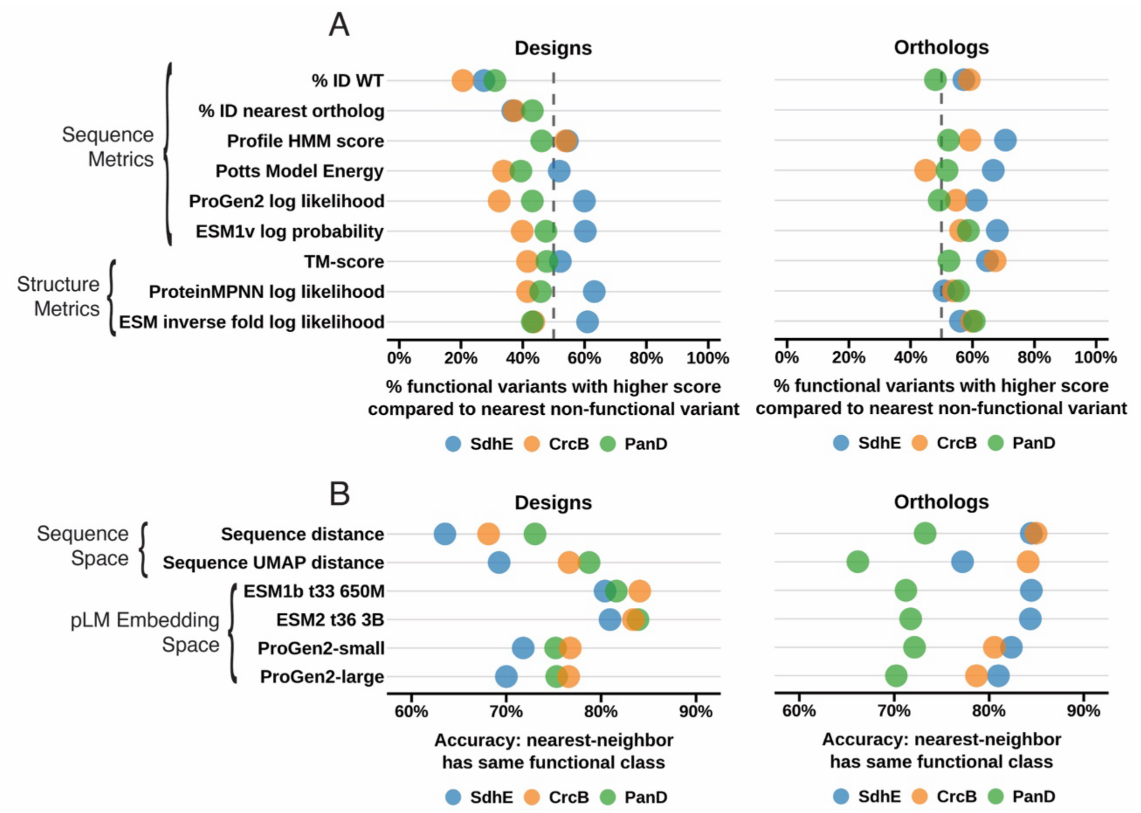


Supplemental Figure 9. Accuracy of function prediction in local sequence space and clustering in pLM embedding space. (A) The percent of functional/non-functional sequence pairs where the functional protein has a more favorable score across all design (left) and ortholog (right) libraries. (B) Accuracy that the nearest neighbor is the same functional class based on sequence distance or cosine-based embedding distance across all design (left) and ortholog (right) libraries.

**Supplemental Table 1. The distribution of functional and non-functional designs relative to orthologs**

| | | | Between orthologs | | | |
|---|---|---|---|---|---|---|
| Gene | Functional | Non-functional | Total | Functional (% between orthologs | Functional (% of all functional) | Non-functional (% of all non-functional) |
| SdhE | 2102 | 3419 | 1474 | 52% | 768 (36%) | 706 (21%) |
| CrcB | 349 | 1442 | 311 | 65% | 202 (58%) | 109 (8%) |
| PanD | 297 | 1611 | 1013 | 26% | 267 (90%) | 746 (46%) |
